# Endocrine fine-tuning of daily locomotor activity patterns under non-starving conditions in Drosophila

**DOI:** 10.1101/2020.02.13.947556

**Authors:** Dennis Pauls, Mareike Selcho, Johanna Räderscheidt, Kelechi M. Amatobi, Agnes Fekete, Markus Krischke, Christiane Hermann-Luibl, Ayten Gizem Ünal, Nadine Ehmann, Pavel M. Itskov, Charlotte Helfrich-Förster, Ronald P. Kühnlein, Martin J. Mueller, Christian Wegener

**Author notes:** **Correspondence**: Dennis Pauls, Department of Animal Physiology, Institute of Biology, Faculty of Life Sciences, Leipzig University, Talstraße 33, 04103 Leipzig, Germany, Christian Wegener, Neurobiology and Genetics, Theodor-Boveri-Institute, Biocenter, University of Würzburg, Am Hubland, 97074 Würzburg, Germany.

## Abstract

Animals need to balance competitive behaviours to maintain internal homeostasis. The underlying mechanisms are complex, but typically involve neuroendocrine signalling. Using *Drosophila*, we systematically manipulated signalling between energy-mobilising endocrine cells producing adipokinetic hormone (AKH), octopaminergic neurons and the energy-storing fat body to assess whether this neuroendocrine axis involved in starvation-induced hyperactivity also balances activity levels under *ad libitum* access to food.

Our results suggest that AKH signals via two divergent pathways that are mutually competitive in terms of activity and rest. AKH increases activity via the octopaminergic system during the day, while it prevents high activity levels during the night by signalling to the fat body. This regulation involves feedback signalling from octopaminergic neurons to AKH-producing cells (APCs). APCs are known to integrate a multitude of metabolic and endocrine signals. Our results add a new facet to the versatile regulatory functions of APCs by showing that their output contributes to shape the daily activity pattern under *ad libitum* access to food.

## 2. Introduction

Foraging behaviour is fundamental to fitness and survival of all animals. When food is scarce, starving animals explore their environment in order to locate food sources. Depending on the amount of available energy and time of the day, foraging can suppress other essential behaviours like sleep. Upon food restriction, rodents and other vertebrate species increase their activity which can be interpreted as foraging [1,2]. In humans, anorexia nervosa is associated with sometimes even excessive hyperactivity, likely triggered by a decreased titre of the peptide hormone leptin [1,3]. Also in the fruit fly, *Drosophila melanogaster*, food deprivation induces locomotor hyperactivity (food search) [4,5], which is triggered by a signalling pathway comprising the peptide hormone adipokinetic hormone (AKH, an analogue to vertebrate glucagon) [4,6–8], and the biogenic amine octopamine (OA) [5,9]. AKH is produced and released by endocrine cells in the glandular part of the corpora cardiaca (CCA; a homologue of the vertebrate anterior pituitary gland and an analogue of mammalian pancreatic alpha cells) [10–12] and is required to maintain haemolymph sugar and lipid levels [4,7,8,11]. Upon starvation, carbohydrate levels in the haemolymph decrease, which is indirectly sensed by the AKH-producing cells (APCs) through ATP-sensitive potassium channels and AMP-activated protein kinase (AMPK) [6,13]. Similar to glucagon, AKH acts on energy stores in the fat body by mobilising carbohydrates in form of the haemolymph sugar trehalose (hypertrehalosaemic effect), and by mobilising lipid depots (adipokinetic effect) [8,14–16].

OA, an analogue to vertebrate noradrenaline, is considered to act as an insect stress or arousal hormone [17]. OA is produced by octopaminergic neurons (OANs) whose cells bodies are located mostly along the midline in the brain and ventral nerve cord [18]. Among other functions, OA exerts myotropic effects on skeletal muscles [19] and mediates the adaptation of muscle metabolism to the physiological demands of locomotion [20–23].

Both AKH and OA are crucial for starvation-induced hyperactivity, as this behaviour is absent in flies with impaired AKH or OA signalling [4,5,7–9]. AKH activates a subset of OANs which are required for starvation-induced hyperactivity [5,9]. These neurons express receptors for AKH and *Drosophila* insulin-like peptides (DILPs). DILPs act antagonistically to AKH, and suppress starvation-induced hyperactivity via the OANs [9]. Yu and colleagues (2016) showed that the AKH-dependent increase in locomotor activity upon starvation represents a direct neuromodulatory and behavioural effect of AKH and is not an indirect consequence of altered energy metabolism [9].

Remarkably, studies in various insect species suggest that AKH and also OA play a role in the regulation of locomotor activity when the insects have free access to food (reviewed in [24]). For example, the circulating haemolymph titres of AKH correlate positively with daily rhythms in walking activity in the linden bug *Pyrrhocoris apterus* [25,26]. In *Pyrrhocoris* [27,28], crickets [29] and the cockroach *Periplaneta americana* [30,31], injection into the haemolymph or topical application of AKH induces a strong increase in locomotor activity in *ad libitum* fed animals, independent of the time of day [28,30]. Also in *Drosophila*, genetic overexpression of AKH enhances locomotor activity when food is freely available [32]. Unlike for starvation-induced hyperactivity, however, it is unclear whether and how AKH:OAN signalling contributes to shape daily activity under *ad libitum* feeding conditions, or whether increased AKH signalling is a consequence of a locomotor-induced increase in energy demands. Cockroach AKHs are known to increase the spontaneous neuronal activity of octopaminergic dorsal unpaired median neurons in the central nervous system, which express a specific G-protein coupled receptor (AKHR) [30,31]. Yet, the functional significance of AKHRs on OANs under *ad libitum* feeding conditions has not been rigorously tested.

We here used the genetically tractable fruit fly to experimentally address the role of AKH:OAN signalling for locomotor activity under non-starved conditions. We find that even under these conditions AKH signalling to OANs participates in shaping diel activity levels. Surprisingly, AKH-induced energy mobilisation appears to antagonize the AKH:OAN-dependent promotion of activity by preventing abnormally high nocturnal activity levels during an initial phase of starvation. In turn, OA modulates the activity of APCs via a feedback signalling comprising OAMBII receptors. Taken together, our results indicate that AKH:OAN and AKH:fat body signalling in *Drosophila* helps to balance and adjust locomotor activity in the ambivalence between foraging and rest.

## 3. Results

To validate our experimental conditions, we first confirmed that AKH is required for starvation-induced hyperactivity and affects starvation resistance (Fig.S1–S2). As previously shown [4,7,8], starved control flies showed a significant starvation-induced hyperactivity compared to fed controls (Fig.S1–S2). This starvation-induced hyperactivity was impaired in flies lacking APCs due to the targeted expression of pre-apoptotic genes *head involution defective* (*hid*) and *reaper* (*rpr*) [33,34] using *Akh-Gal4* [4]. Increased lipid stores in fat body cells contribute to the higher starvation resistance observed in flies lacking AKH signalling [8,15,16]. In line with this, *Akh>hid,rpr* flies showed higher starvation resistance during activity recording, as on day 3 more than 80% of flies were still alive, compared to less than 15% of starved control flies (Fig.S3).

### Octopaminergic neurons modulate diurnal activity levels in fed flies

In addition to AKH, OA signalling is essential for starvation-induced hyperactivity [5]. We first tested whether the OA neuronal system is also involved in modulating locomotor activity in flies that have permanent access to food. To manipulate OANs, we used the *Tdc2-Gal4* line. Tdc2 (Tyrosine decarboxylase 2) is the enzyme necessary for the rate-limiting decarboxylation of tyrosine to tyramine (TA) which then is further hydroxylated by tyramine beta-hydroxylase (TßH) to OA [35]. *Tdc2-Gal4* specifically marks OANs and tyraminergic neurons [36–39]. We first tested whether *Tdc2-Gal4*-positive neurons (Tdc2Ns) are involved in the modulation of locomotor activity in fed flies and ablated Tdc2Ns by the expression of UAS-*hid,rpr*. As expected, Tdc2>hid,rpr flies showed significantly reduced locomotor activity during the day, while activity levels during the night were unaffected (Fig.1A-C). We next blocked neuronal transmission of Tdc2Ns using UAS-ΔOrk [40] and UAS-*shibire^ts^* [41] to exclude any side effects on activity due to cell ablation. In line with the previous results, both electrical and chemical silencing of synaptic transmission reduced locomotor activity levels specifically during the day (Fig.1D-I).

**Figure 1:**
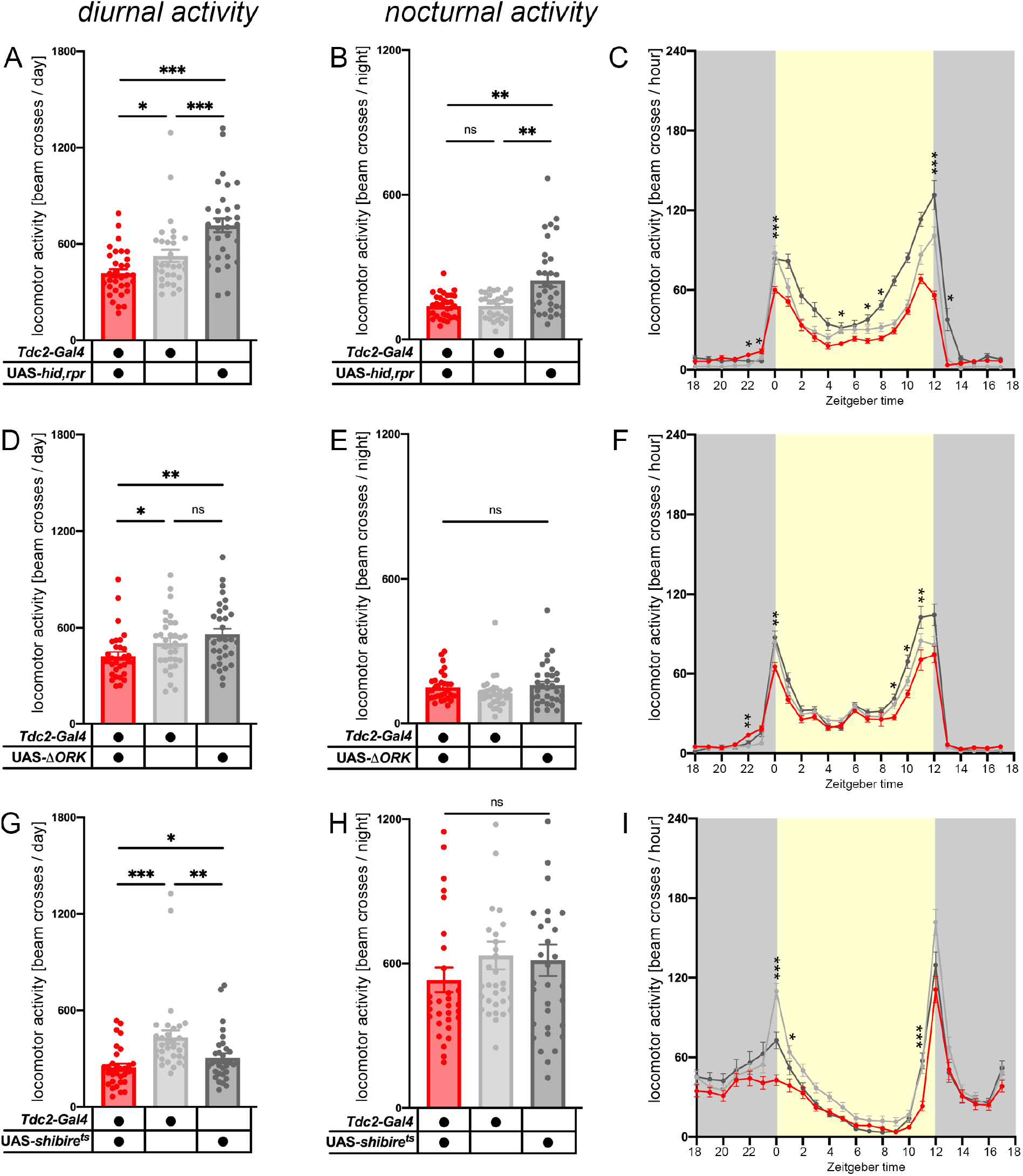
Tdc2-Gal4-positive neurons modulate activity during the day under non-starvation conditions. **(A-C)** Activity levels are reduced during the day in Tdc2>hid,rpr flies compared to genetic controls (p<0.05 to Gal4/+ and p<0.001 to UAS/+). **(D-F)** Neuronal silencing of Tdc2Ns (Tdc2-Gal4-positive neurons) revealed a similar effect with reduced activity of experimental flies during the day. **(G-I)** Conditional thermogenetic silencing of Tdc2Ns using UAS-shi^ts1^ at restrictive temperature also reduced activity levels during the day (p<0.001 to Gal4/+ and p<0.01 to UAS/+). n=26-32, significance levels: p>0.05 ns, p<0.05 *, p<0.01 **, p<0.001 ***. Error bars indicate SEM.

Our results are consistent with an earlier study showing that *Tdc2^R054^* and *TβH^nm18^* mutants increase sleep mainly during the day [42,43], but partially differ from results published by Crocker and Sehgal: here Kir2.1-mediated electrical silencing of *Tdc2Ns* consistently decreased locomotor activity, but also increased night sleep [42]. Nonetheless, the results show that OA/TA signalling promotes locomotor activity and hence wakefulness during the night [42,44] or under starvation [5,9], and in addition shapes the general locomotor activity level during the day when flies have permanent access to food.

### AKH:OAN signalling is important for the modulation of day-time activity

Our results and data from *Tdc2* mutants [42] showed that impaired OA signalling reduced locomotor activity specifically during the day. We thus hypothesized that OANs signal primarily during the day but not during the night to modulate locomotor activity under *ad libitum* feeding conditions. This is consistent with the view of OA (and TA) being a stress hormone that is released during physical activity [17,23]. AKH positively modulates OANs in cockroaches and fruit flies [9,30], which predicts that ablation of APCs and down-regulation of the AKH receptor (AKHR) specifically in Tdc2Ns will result in reduced locomotor activity during the day (when OA signalling is effective), but unaltered locomotor activity during the night (when OA signalling is not present under *ad libitum* feeding conditions). Indeed, similar to the ablation of Tdc2Ns (Fig.1A-C), *Akh>hid,rpr* and *Tdc2>AkhR-RNAi* flies showed significant reduced locomotor activity during the day, while locomotor activity was unaffected during the night (Fig.2A-F). In order to demonstrate that the effect of the AKHR knockdown in Tdc2Ns was specific to day-time locomotor activity and not due to a general impairment of motor capability, we performed a startle-induced negative geotaxis (SING) climbing assay [45]. The performance of *Tdc2>AkhR-RNAi* flies was indistinguishable from genetic controls in this climbing assay, suggesting that motor capability was not generally affected by our genetic manipulation (Fig.S4). Interestingly, while the ablation of APCs led to a reduction in activity throughout day-time (Fig.2A-C), the average activity profile in *Tdc2>AkhR-RNAi* flies suggests that the reduction in activity is mainly based on a reduced activity immediately after the morning activity bout, while locomotor activity around the evening peak was indistinguishable from genetic controls (Fig.2F). This day-time specific reduction of locomotor activity was partially rescued when we restricted the expression of *AkhR-RNAi* to Tdc2Ns in the brain, using *tshGal80* (Fig.S5) [46]. Expression of GFP in *Tdc2,tshGal80>10xmyr::GFP* flies revealed a strong reduction of labelled Tdc2Ns in the ventral nerve cord (VNC). From ~39 *Tdc2-Gal4-* positive VNC neurons [47] ~7 cells (7.4 ± 2.07, n=5) escaped *tshGal80*-dependent suppression (Fig.S6), while the number of Tdc2Ns in the brain was not affected. These results suggest that Tdc2Ns in the VNC are involved in the AKH:OAN signalling. To confirm the expression of AKHR in cells in the VNC we performed RT-PCR. We found AKHR to be expressed in isolated adult VNCs, as well as in adult brains and the larval fat body (Fig.S7).

**Figure 2:**
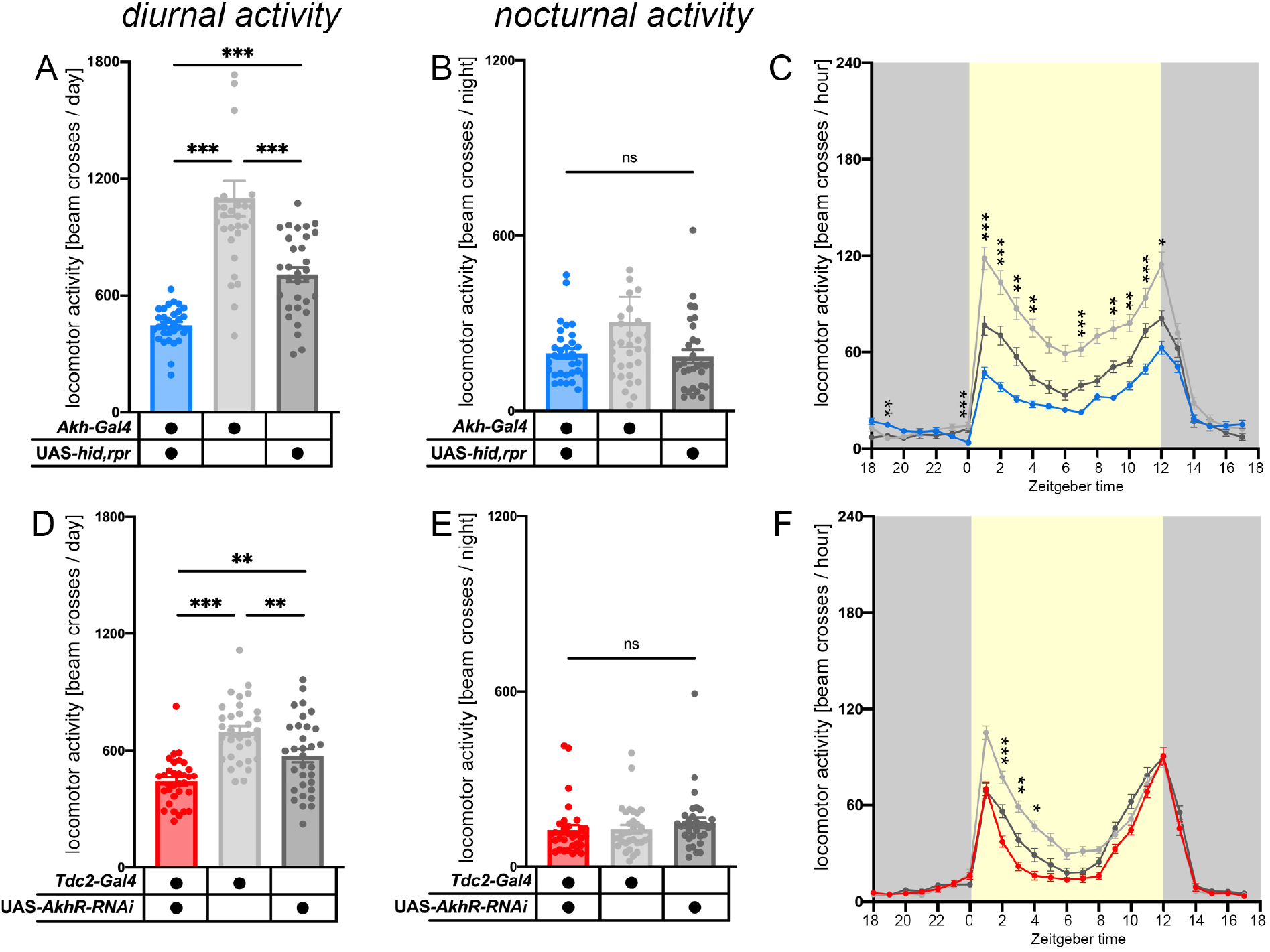
AKH:OAN signalling is important for the modulation of day-time activity. **(A-C)** Akh>hid,rpr flies showed significantly reduced activity (all p<0.001) specifically during the day. **(D-F)** Knockdown of AKHR in Tdc2Ns (Tdc2-Gal4-positive neurons) specifically affected locomotor activity during the day (p<0.01 to Gal4/+ and p<0.001 to UAS/+). n=26-32, significance levels: p>0.05 ns, p<0.05 *, p<0.01 **, p<0.001 ***. Error bars indicate SEM.

The results so far are consistent with data from the cockroach *Periplaneta americana*, where AKHR is expressed in thoracic and abdominal octopaminergic dorsal unpaired median neurons and signals both via G_q_ and G_s_ to activate cAMP and Ca^2+^-dependent second messenger pathways [30,31]. To confirm AKH signalling to OANs located in the VNC, we bath-applied AKH to isolated VNCs (detached from the brain) and recorded changes in intracellular cAMP or free Ca^2+^ levels, respectively, in Tdc2Ns expressing the genetically encoded Epac1-camps50A [48] or GCaMP6m [49] sensors, focusing on the mesothoracic cluster.

All mesothoracic Tdc2Ns responded to the synthetic adenylate cyclase agonist NKH477 or the nicotinic receptor agonist carbamylcholine with robust increase in cAMP/free Ca^2+^ levels showing that neurons were functional in our preparations. Upon bath-application of 10^-5^M AKH, Tdc2Ns in the mesothoracic cluster responded with a significant cAMP increase, while application of haemolymph-like HL3.1 saline solution as vehicle control had no effect (Fig.S8) [50]. In contrast, free intracellular Ca^2+^ levels did not significantly change in Tdc2Ns in the VNC upon AKH bath application compared to HL3.1 vehicle control (Fig.S10). These results suggest that activated AKHR modulate OAN activity via cAMP-dependent pathways. Of note, the observed AKH-induced increase in cAMP levels occurred with a time lag of at least 15 minutes after bath application of the peptide (Fig.S8). We next overexpressed AKHR in Tdc2Ns and repeated cAMP imaging in the mesothoracic cluster. Overexpression of AKHR did not significantly increase cAMP beyond the level observed in controls (Fig.S9). Although the response to exogenous AKH occurred significantly faster, responses were still delayed with a mean lag time of around 10 minutes (Fig.S9). This suggests that AKH induces slow cAMP elevations in OANs even if AKHR is overexpressed in these neurons, potentially in a dose-dependent manner.

To provide further genetic evidence for the modulatory role of AKHR in OANs, we rescued AKHR expression specifically in Tdc2Ns in an *AKHR^1^* mutant background and monitored locomotor activity (Fig.3A-C) [15]. Rescue flies showed significantly increased locomotor activity during the usual activity bouts in the morning and evening and during the early and late night, compared to AKHR-deficient controls (Fig. 3A-C). No difference in locomotor activity was apparent during the midday siesta, which is reflected by the lack of significant day-time activity differences compared to genetic controls.

**Figure 3:**
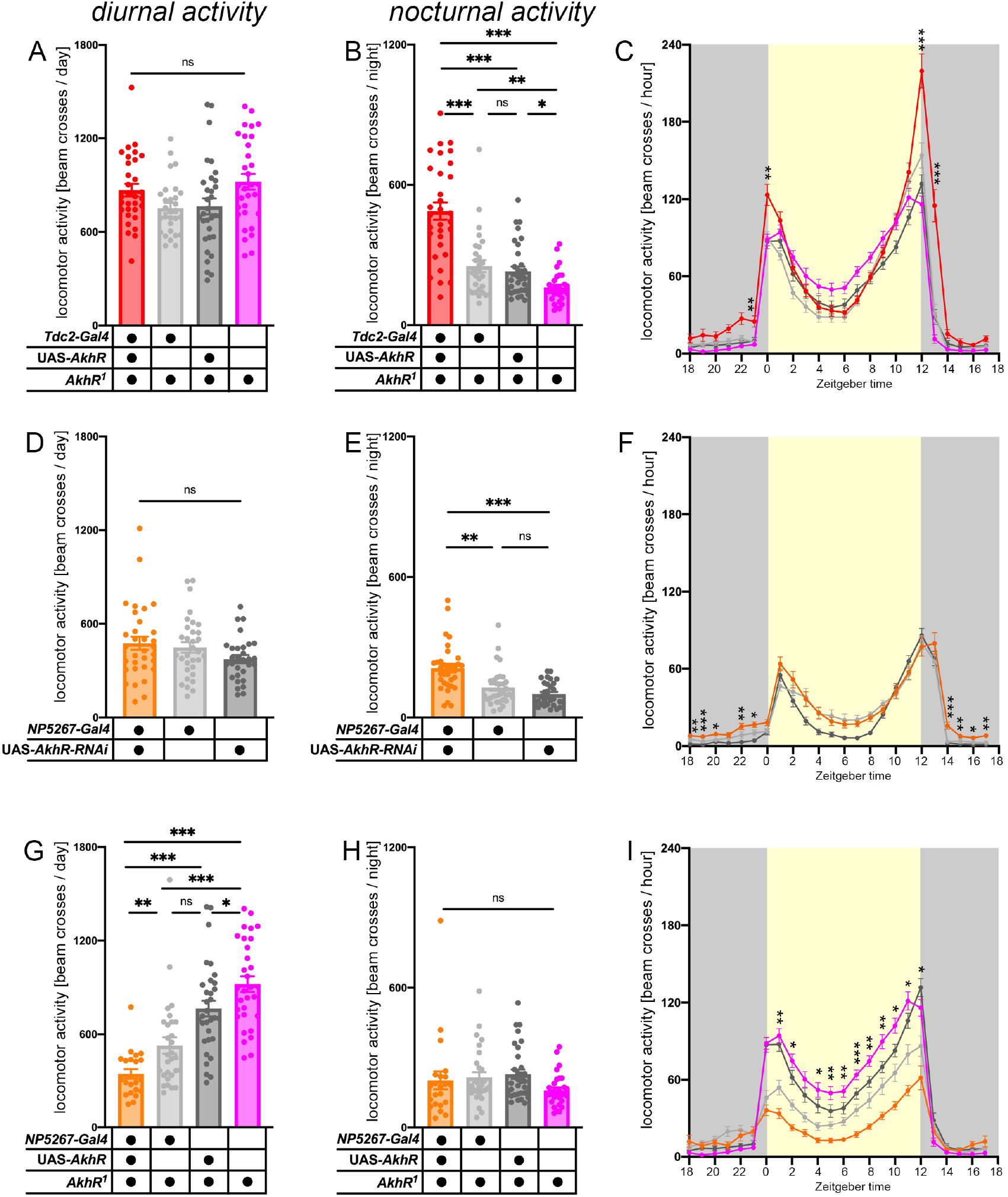
AKH balances day and night locomotor activity via different signalling routes. **(A-C)** Genetic rescue of AKHR in Tdc2Ns (Tdc2-Gal4-positive neurons) rescued locomotor activity during the day (all p>0.05); however, AkhR^1^,Tdc2>AkhR,AkhR^1^ flies showed significantly higher activity during morning and evening activity bouts (all p<0.001 at ZT0 and ZT12). In addition, experimental flies showed significantly increased activity levels during the night (all p<0.001). **(D-F)** Flies with a fat body-specific knockdown of AKHR showed increased levels of activity during the night (p<0.01 to Gal4/+ and p<0.001 to UAS/+. **(G-I)** Specific rescue of AKHR in the fat body elicited the opposite effect seen to the rescue in OANs (see A-C). Expression of AKHR specifically in the fat body rescued activity levels during the night; in contrast, locomotor activity was significantly reduced during the day (all p<0.01). Note that the experiments in A-C and G-I were conducted at the same time. The controls shown are identical in the two experiments. n=13-32 for behavioural data, significance levels: p>0.05 ns, p<0.05 *, p<0.01 **, p<0.001 ***. Error bars indicate SEM.

Taken together, the results so far suggest that AKH modulates OANs in the VNC (and potentially in the brain) [5,9] to increase locomotor activity during the day when flies have permanent access to food. Yet, the AKHR is most strongly expressed by the fat body [15,24] where it mediates the hypertrehalosaemic effect of AKH and mobilizes stored energy in form of the haemolymph sugar trehalose (Fig.7). To test whether AKH:fat body signalling affects locomotor activity under *ad libitum* feeding conditions, we first knocked down AKHR expression specifically in the fat body using *NP5267-Gal4. NP5267>AkhR-RNAi flies* showed significantly increased locomotor activity during the night (Fig.3D-F). Further, specific rescue of AKHR in the fat body in a *AkhR^1^-* mutant background significantly reduced locomotor activity during morning and evening activity bouts as well as during midday siesta (phenocopying the impairment of AKH:OAN signalling; see Fig.1 and Fig.2), while locomotor activity appeared to be normal during the night (Fig.3G-I).

### OAN signalling modulates AKH cells in the corpora cardiaca

Our results so far support a role for AKH:OAN signalling under *ad libitum* feeding conditions. We next wondered whether OANs in turn provide feedback signals to the APCs in the corpora cardiaca (CCA). Thus, we traced the projections of Tdc2-positive cells in whole body sections of flies using a Tdc2 antibody (Fig.4A-B). Besides arborizations in the brain and VNC, we found Tdc2-positive fibres projecting to the CCA-hypocerebral ganglion complex (CCA-HG complex, also known as stomodaeal ganglion; Fig.4B, inlet) presumably via the stomodaeal nerve (Fig.4B, asterisk). Thus, we assumed that OA may be released in vicinity of the AKH cells. Among the different OA receptors, OAMBII has been responsible for most OA actions [17], and increases intracellular Ca^2+^ levels in heterologous expression systems as well as in *Drosophila* neurons [51–53]. Using primers targeting different exons of the OAMBII gene [54], we confirmed the expression of the OAMB-K3 isoform (Fig.4D, OAMB-K3 contains exon 7 but not 8) in the CCA-HG complex, while signals for the OAMB-AS isoform (OAMB-AS contains exon 8 instead of 7) were not detected (Fig.4C). To test whether APCs express functional OA receptors, we expressed the genetically encoded Ca^2+^ sensor GCaMP6m in APCs and imaged the effect of bath-applied OA. For these experiments, we used the whole larval central nervous system with attached ring glands as it was not feasible to dissect intact adult CCA-HG-brain preparations. Since at low trehalose levels the Ca^2+^ concentration is generally high in APCs, we adopted a procedure developed by Kim and Rulifson (2004) to image Ca^2+^ changes in these cells: preparations were first pre-incubated in 80mM trehalose solved in HL3.1 which decreased free intracellular Ca^2+^ concentration to basal level (Fig.4E: arrow No.1 at 100s). Subsequent application of 1mM OA + 80mM trehalose (Fig.4E: arrow No.2 at 300s) significantly increased intracellular Ca^2+^ levels indicated by an increase of fluorescence intensity (Fig.4E, blue line). In contrast, no changes in Ca^2+^ levels were visible after bath application of the control solution (Fig.4E, grey line), suggesting that APCs respond with increased intracellular Ca^2+^ levels to OA (Fig.4F).

**Figure 4:**
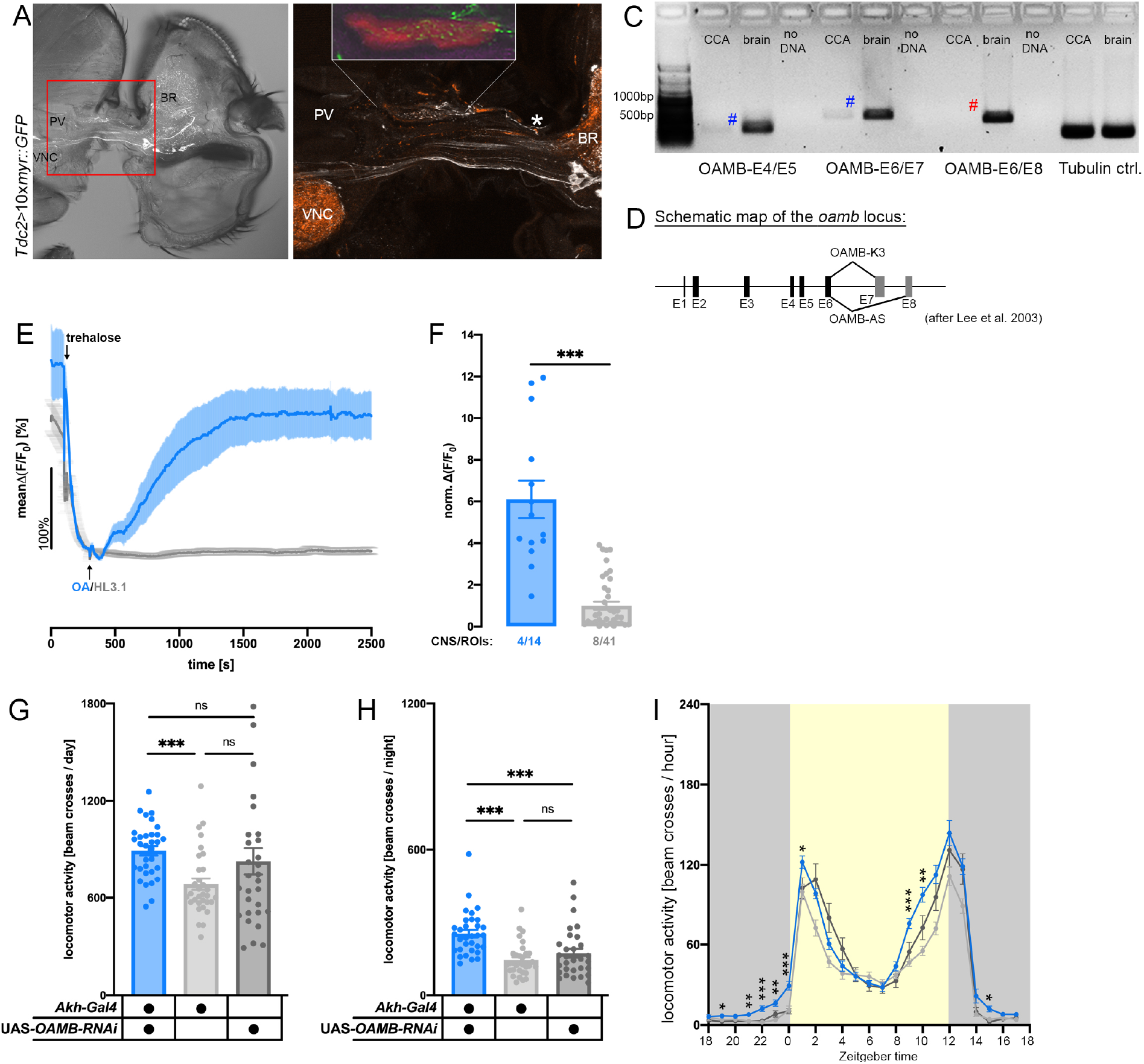
AKH release is modulated by OAN feedback signalling. **(A-B)** Whole body sections of flies displayed arborizations of Tdc2-positive neurons (white) in the CCA-HG complex (inlet in (B)). (B) shows a magnification of the red squared region in (A). The CCA-HG complex is highlighted in red in the inlet. White: anti-Tdc2; orange: anti-Synapsin. **(C-D)** RT-PCR analysis confirmed that the OA receptor OAMBII-K3 isoform is expressed in the CCA-HG complex (blue #), but not the OAMBII-AS isoform (red #). **(E-F)** Average changes in Ca^2+^ levels of larval AKH cells to bath applied OA and HL3.1, respectively (Akh>GCaMP6m). AKH cells were pre-incubated with 80mM trehalose to reduce intracellular Ca^2+^ levels to baseline (arrow at 100s). AKH cells responded to bath applied OA (arrow at 300s) with a strong increase in intracellular Ca^2+^ levels (blue line; (F)); in contrast, no significant responses were obvious in response to the vehicle control (grey line). **(G-I)** Akh>OAMBII-RNAi flies showed significantly increased activity during the night (all p<0.001). n=29-32 for behavioural data, significance levels: p>0.05 ns, p<0.05 *, p<0.01 **, p<0.001 ***. [CCA= corpora cardiaca, Ctrl= control, BR=brain, E=exon, HG=hypocerebral ganglion, OA=octopamine, PV=proventriculus, ROI=region of interest, VNC=ventral nerve cord]. Error bars indicate SEM.

As our results suggest that OANs signal to APCs via the OAMBII receptor, we next performed a knockdown of OAMBII in these cells (*Akh>OAMBII-RNAi*). The knockdown resulted in significantly increased nocturnal activity (Fig.4G-I), suggesting that OANs modulate AKH release. Modulation of APC activity by OA is likely a common feature among insects, as OA was found to modulate AKH secretion in the locust [55,56]. In locusts, the CCA is not innervated by OANs, and OA likely acts in a hormonal fashion [57]. In *Drosophila*, however, *Tdc2*-positive fibres innervate the CCA-HG complex (Fig. 4A,B) and OA may act in a paracrine fashion.

### AKH:fat body signalling prevents an increase of activity during the night

Our results suggest that - opposite to the stimulatory locomotor effect of AKH:OAN signalling - AKH:fat body signalling prevents an increase of activity with the most pronounced effect during the night. Previous studies suggested that haemolymph trehalose levels are at a trough at the end of the night and immediately after lights on [58]. We therefore hypothesized that AKH modulates locomotor activity in a balanced state-dependent manner. AKH:OAN signalling may promote activity during the day, while AKH:fat body signalling stabilises rest during the night.

To test this hypothesis, we monitored locomotor activity of *NP5267>AkhR-RNAi* for three days, but this time under starved conditions. We assumed that flies lacking AKH:fat body signalling cannot compensate for the lack of food with energy mobilization from the fat body and thus will show an earlier onset of nocturnal starvation-induced hyperactivity than control flies. Flies were recorded from ZT0 (lights-on; ZT=Zeitgeber time) in tubes that contained agarose providing water to prevent dehydration, but lacked sugar or other nutrients. All flies showed starvation-induced hyperactivity during the experiment (Fig.5A-C). For *NP5267>AkhR-RNAi* flies, however, this hyperactivity started earlier and became already obvious during the first night around 12h after starvation onset (Fig.5A; red arrow). In contrast, control flies became hyperactive not until the second day (Fig.5B-C; red arrows), with starvation-induced hyperactivity fully developed first during the second night in both genetic controls (Fig.5D-F). Thus, while *NP5267>AkhR-RNAi* flies were significantly hyperactive from the first night on in response to starvation, endogenous AKH:fat body signalling prevented undue high nocturnal activity levels in consequence of food search in control flies during the first night, presumably by mobilizing energy from the fat body into the haemolymph (Fig.5D-F).

**Figure 5:**
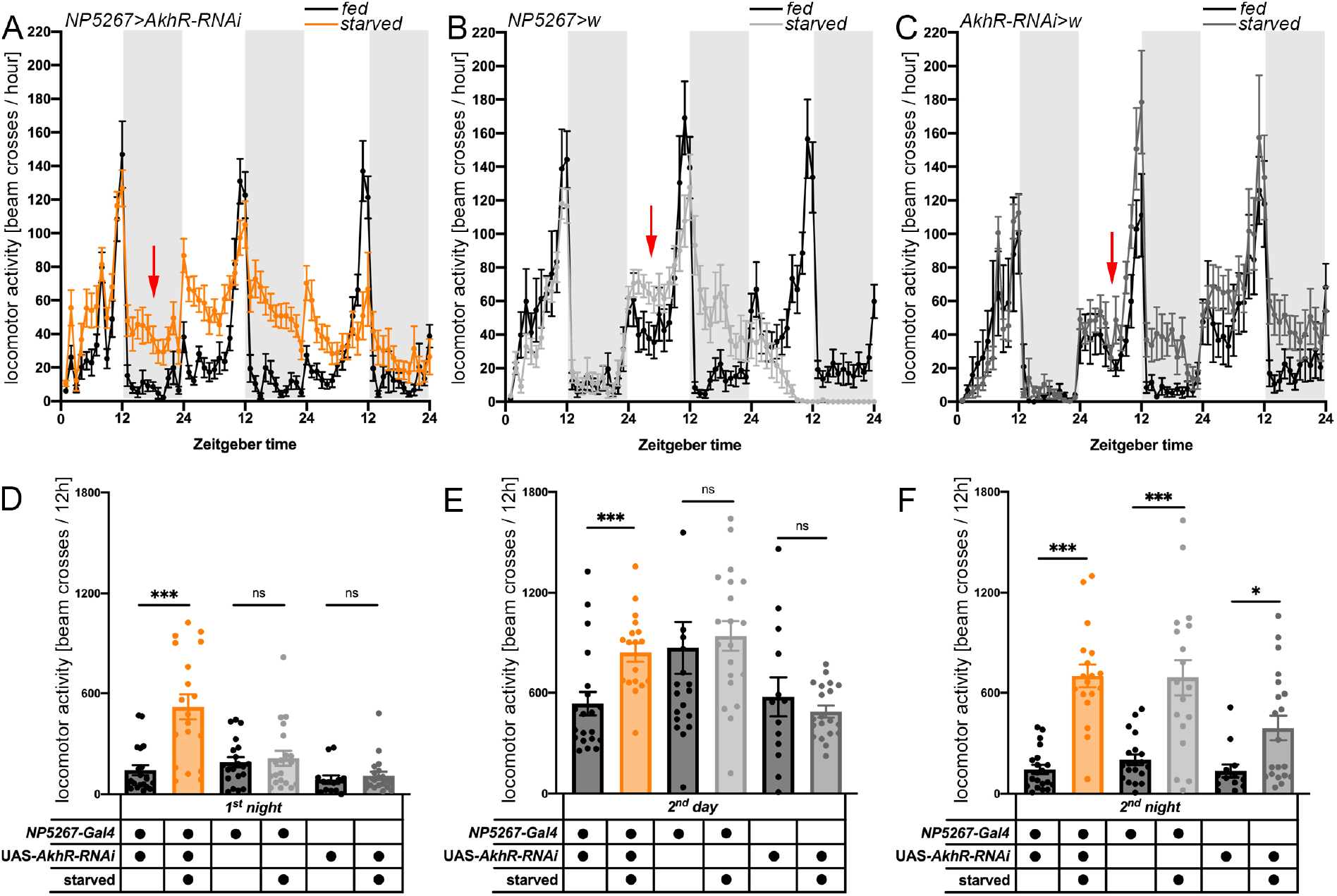
Onset of starvation-induced hyperactivity. **(A-F)** NP5267>AkhR-RNAi flies showed an early onset of starvation-induced hyperactivity during the first night (after ~12h of starvation; red arrow). In contrast, AKH:fat body signalling in control flies allowed a short-term adaptation under starvation and prevented abnormally high night activity. Red arrows indicate first obvious appearance of starvation-induced hyperactivity, n=13-32; significance levels: p>0.05 ns, p<0.05 *, p<0.01 **, p<0.001 ***. Error bars indicate SEM.

### Impaired AKH-dependent carbohydrate mobilisation is compensated by dietary uptake

So far, our results suggest that AKH signalling modulates locomotor activity not only under starved conditions, but also when animals have free and permanent access to food. The effect was most obvious during the night when feeding-related behaviours are normally absent or strongly reduced [58]. We assumed that during these nocturnal non-feeding phases, flies usually compensate the reduced dietary carbohydrate intake via the hypertrehalosemic effect of AKH. However, flies deficient of AKH:fat body signalling are unable to maintain high trehalose levels [4,7,11]. This may lead to increased AKH release, which in turn will promote foraging activity via the integrated increase of AKH:OAN-signalling (Fig.5A) especially towards the end of the night, analogous to but less pronounced as starvation-induced hyperactivity. To find out whether APCs of flies with impaired AKH signalling are more active during the night, we first used calcium-dependent nuclear import of LexA (CaLexA) [59], an integrator of long-term Ca^2+^ activity. As expected, CaLexA-driven GFP expression increased significantly upon 24 hours of starvation in *AKH>CaLexA* flies (Fig.S11B-C, F-G), showing that the method is apt to monitor APC activity.

On food, we found a weak and statistically insignificant increase in CaLexA-dependent GFP expression in *Akh>CaLexA* controls between end of the day (ZT10) and middle (ZT18) or end (ZT22) of the night (Fig.S11E, F-G). In *Akh;amon-RNAi>CaLexA* flies with reduced AKH signalling [11], we found a slight but insignificant increase in CaLexA-driven GFP expression compared to controls (Fig. S11D, F-G). These results suggest that APCs generally have a low activity on food, and speak against a significantly increased need for energy mobilization in flies with impaired AKH signalling under our experimental conditions. We therefore tested whether the loss of AKH-dependent carbohydrate mobilization is compensated by dietary carbohydrate uptake. First, we quantified haemolymph sugar levels during the light phase under starved and *ad libitum* feeding conditions using Ultra performance liquid chromatography - tandem mass spectrometry. Indeed, APC-ablated flies under fed conditions showed similar haemolymph levels of sucrose, fructose, trehalose and glucose, although trehalose and fructose levels were somewhat reduced (Fig.6A-D). Under starved conditions, however, trehalose levels became depleted in APC-ablated flies (Fig.6C), as previously shown [7]. Also, glucose levels were strongly though not significantly reduced (Fig.6D). In contrast, fructose and sucrose levels did not significantly differ compared to controls and were generally very low (Fig.6A-B), likely because the haemolymph titres of theses sugars originate mostly from food uptake. These results fully support our assumption that flies deficient in AKH signalling compensate impaired carbohydrate mobilisation by food uptake, and explain the similar activity levels of APCs between controls and AKH down-regulated flies as determined with CaLexA.

**Figure 6:**
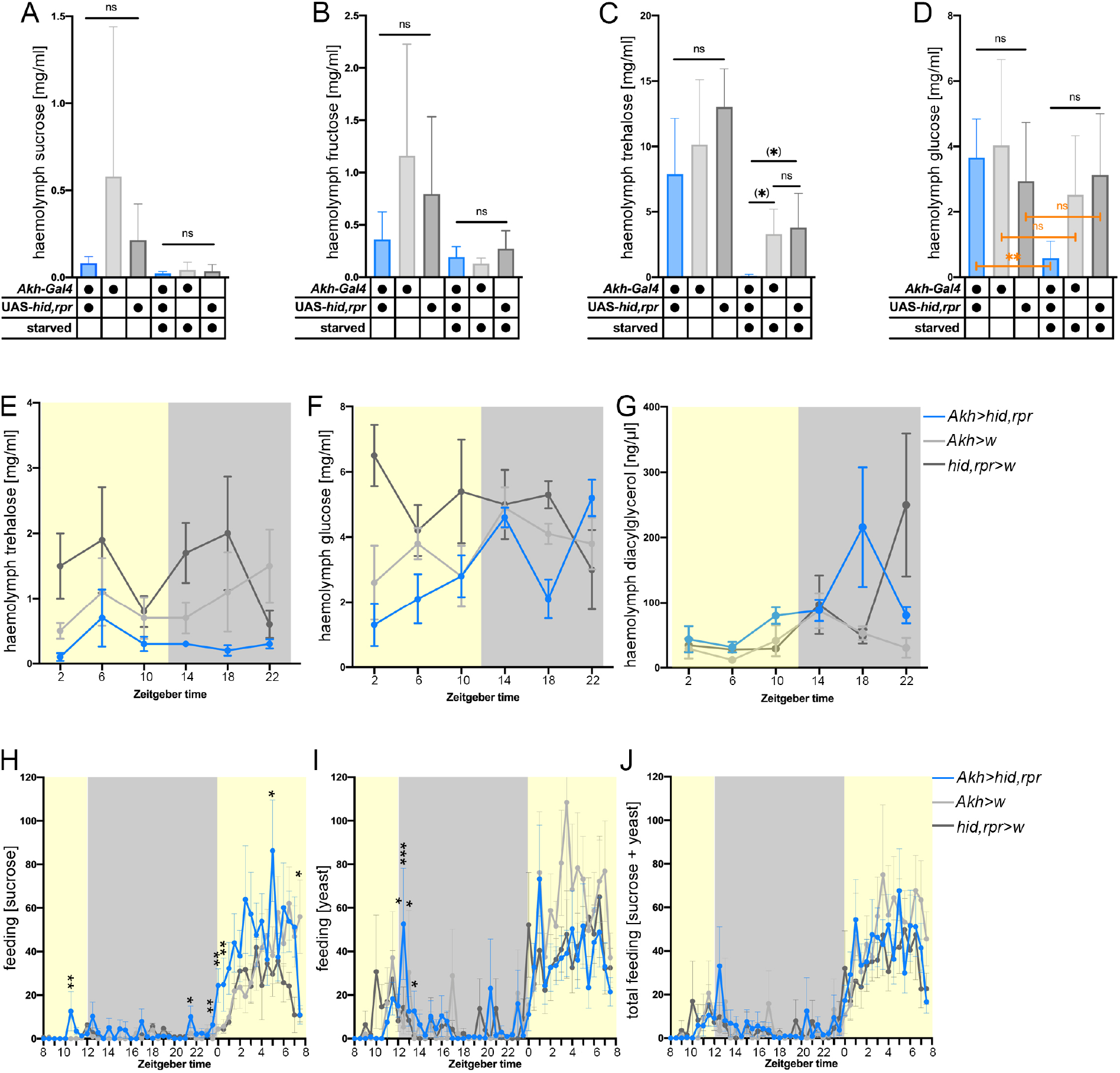
Carbohydrate haemolymph titres and feeding activity. **(A-D)** Haemolymph sugar levels analyzed by UPLC-MS/MS in Akh>hid,rpr flies and controls: while sucrose and fructose levels did not differ between genotypes and conditions (starved vs. fed), trehalose levels were depleted in experimental flies under starvation conditions (p=0.078 to Gal4/+ and p=0.063 to UAS/+). In addition, glucose levels were reduced in Akh>hid,rpr flies between fed and starved conditions (p<0.01), without any effect in control flies (p>0.05; all N=3-4). **(E)** In control flies, haemolymph trehalose levels rise during the night. In contrast, trehalose levels stay low during the night in NP5267>AkhR-RNAi flies. **(F)** Haemolymph glucose levels increase throughout 24h in experimental flies indicating higher food intake compared to controls. **(G)** In contrast to haemolymph sugar levels, no obvious change was detectable on haemolymph lipid levels. **(H-J)** Time course of feeding (number of sips per 30 min) for sucrose **(H)**, yeast extract **(I)** or sucrose and yeast combined **(J)** in Akh>hid,rpr flies and controls. Significant differences between APC-ablated flies and controls are visible for sucrose feeding mostly at the end of the night and the first hours after lights on, while feeding on yeast or total food consumption was not significantly altered. n=35-36 for feeding data, significance levels: p>0.05 ns, p<0.05 *, p<0.01 **, p<0.001 ***, (p<0.06 (*)). [UPLC-MS/MS=Ultra performance liquid chromatography - tandem mass spectrometry]. Error bars indicate SEM.

Next, we quantified trehalose, glucose, and diacylglycerol levels every four hours during the day under *ad libitum* feeding conditions. In line with the results in Fig.6C, trehalose levels of *Akh>hid,rpr* flies were lower but not significantly reduced compared to both controls (Fig.6E). Again, as expected from previous measurements (Fig.6D), the glucose concentration were similar in both *Akh>hid,rpr* and control flies (Fig.6F).

Yet, the temporal pattern differed. *Akh>hid,rpr* flies showed constant low levels of trehalose during the night, while control flies displayed increased levels of trehalose during rest phases (the siesta and night) suggesting trehalose mobilization from the fat body (Fig.6E). The glucose level stayed constant within a certain range in controls, with a drop at the end of the night in *Akh>w* control flies (Fig.6F). In *Akh>hid,rpr* flies, however, the glucose titre was low at ZT2 and steadily increased until ZT14 to then drop at ZT18 and increase again to a high level at ZT22. This suggests feeding activity in *Akh>hid,rpr* flies especially around ZT22, given that *Akh>hid,rpr* flies are impaired in mobilizing carbohydrates. Interestingly, no obvious differences in haemolymph diacylglycerol levels were detectable between the genotypes (Fig.6G). Strong increases of diacylglycerol occurred at times when the glucose titre dropped even in *Akh>hid,rpr* flies (Fig.6F), providing further evidence for an AKH-independent lipid mobilisation pathway [15].

To directly address food uptake, we used an automated two-choice flyPAD assay [60] and monitored feeding in individual flies on 5mM sucrose vs. 10% yeast (Fig.6H-J, S12). Generally, flies consumed significantly more sucrose and yeast during the day than during the night (Fig.S12). Though not significantly different, sucrose consumption was by trend elevated in *Akh>hid,rpr* flies both during day and night. Analysis at a higher temporal resolution revealed significantly increased sucrose feeding in *Akh>hid,rpr* flies at the end of the night and after lights-on during morning activity compared to controls (Fig.6H). A less pronounced increase in feeding was also visible around the end of the day (coinciding with the start of evening activity), as well as during the night though levels did not reach significance except for ZT 10-11 (Fig.6H). Thus, genetic ablation of APCs led to an earlier onset of feeding activity with a slight general increase of feeding throughout the night. In contrast, total yeast consumption was not significantly affected in *Akh>hid,rpr* flies during the day and night (Fig.S12). Unlike for sucrose feeding, a consistently and significantly elevated level of yeast feeding in *Akh>hid,rpr* flies was also not visible at higher temporal resolution across the day (Fig.6I). Likewise, total feeding was not affected by APC ablation (Fig.6J). These results show that ablation of APCs exerts a specific effect on sucrose but not yeast feeding under our conditions. The observed time course of sucrose feeding in *Akh>hid,rpr* flies (Fig.6H) is compatible with their glucose haemolymph profile (Fig.6F) which shows increasing concentrations during the day, and a drop in the middle of the night when feeding activity was basically absent in *Akh>hid,rpr* flies (Fig.6H).

Taken together, our results propose that *Akh>hid,rpr* flies are not able to mobilize trehalose from the fat body and rescue haemolymph carbohydrate titres by increased food uptake at certain times of the day.

## 4. Discussion

The circadian activity and rest pattern of animals is determined by a multitude of intertwined systems and mechanisms at different levels. The general organisation is in large set out by the circadian clock and the sleep homeostat [61]. Yet, internal and external signals can dramatically alter activity/rest cycles and induce locomotor activity to meet physiological demands or to escape from harmful situations. A particularly well studied example is the starvation-induced locomotor activity in *Drosophila*, which is triggered by the action of the peptide hormone AKH on neurons that release the arousal signal OA [4,5,9]. On food, AKH signalling was thought to not affect general activity levels, as AKH or AKHR mutants show similar activity levels as controls [8,62], and display normal circadian activity patterns [4,63]. Our study now suggests that the unaltered activity levels and activity/rest patterns in AKH and AKHR mutants is an outcome of the combined loss of both branches of AKH signalling to OANs (regulating locomotor activity via feedforward and feedback signalling) and to the fat body (regulating energy mobilisation). By individually manipulating either of the two AKH signalling branches, we were able to uncover that AKH signalling balances and fine-tunes day and night locomotor activity levels when flies have unrestricted access to food.

When AKH:OAN signalling was genetically impaired, flies were significantly less active during the day. During the night, activity levels remained unaffected. In contrast, when AKH:fat body signalling was impaired, the opposite was observed. Activity remained unaffected during the day, but flies were more active during the night. Concomitantly,

*AkhR^1^* mutant flies with rescued AKH:OAN signalling were significantly more active during the night, while day activity was unaffected. Again, the opposite was observed when selectively AKH:fat body signalling was restored in *AkhR^1^* mutants: flies were less active during the day, while normal activity levels were present during the night. Based on our results and available literature data, we propose the following model for the modulatory action of AKH on balancing activity under given conditions (Fig.7). When the fly needs to mobilize energy, for example due to increased physical activity during the day, AKH is released into the circulation to mobilize carbohydrate and lipid fat body stores, and simultaneously activates OANs in the CNS (see Fig.2) [9,64]. The resulting increase in haemolymph trehalose and glucose titres negatively feeds-back to APCs, reducing their secretory activity via opening of ATP-sensitive potassium channels [6,65]. This reduces AKH:fat body signalling, as long as fat body energy stores are not depleted (under fed conditions), and simultaneously prevents a strong activity-promoting effect of time-delayed AKH:OAN signalling. Under starved conditions, however, energy stores become depleted, which leads to a strong and long-lasting increase in the haemolymph AKH titre once carbohydrate levels fall, which in turn stimulates hyperactivity via the activation of OANs [4,7–9].

**Figure 7:**
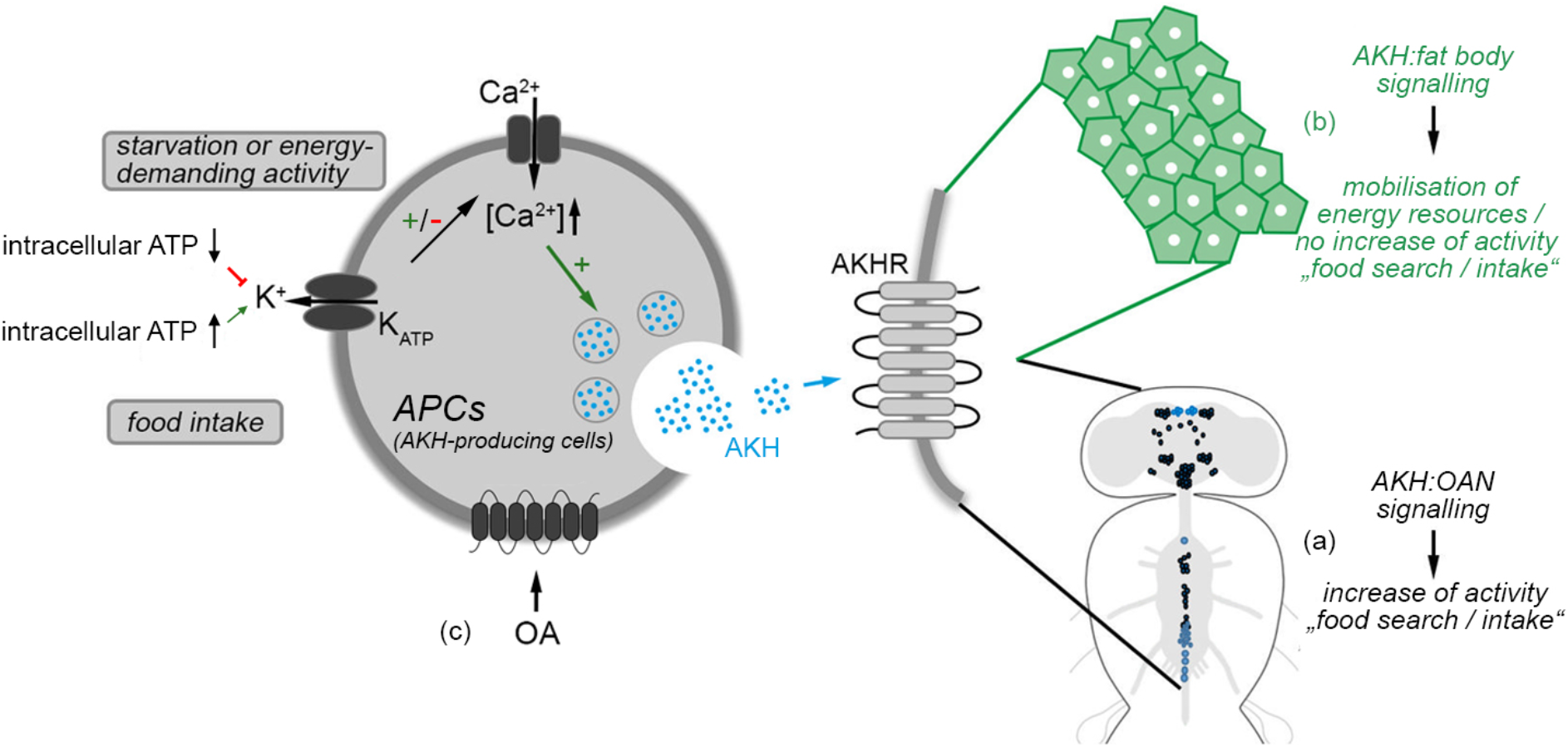
AKH signalling to OANs and the fat body balance competing rest and feeding drives. In response to a negative balance in haemolymph carbohydrate homeostasis (e.g. under starved conditions), AKH is released from neuroendocrine cells in the corpora cardiaca and functions via two mutually competitive signalling pathways: (a) AKH signalling to OANs induces food-search behaviour in response to starvation (during the day); (b) at the same time AKH mobilizes energy from the fat body to restore haemolymph sugar levels; (c) APC activity is regulated by OA-feedback signaling. This will inhibit the activation of locomotor activity by OANs and thereby prevents abnormally high activity levels during the night. As a stress hormone, the release of AKH presumably balances not only locomotor activity to feeding; AKH release will rather induce a general hypermetabolic response, an increase in respiration and immune response [57]. [AKH: adipokinetic hormone; AKHR: AKH receptor; ATP: adenosine triphosphate; OA: octopamine; OAN: octopaminergic neuron]

When AKH:fat body signalling is genetically impaired, flies cannot efficiently mobilise fat body energy stores when haemolymph carbohydrate levels are low. In that case, during the day, haemolymph sugar levels are restored by food uptake, as suggested by the observed increase in trehalose and glucose titres during the light phase and the increased sucrose feeding during morning activity, the main time of food intake [58]. As a consequence, impairing AKH:fat body signalling in flies with free access to food does not induce hyperactivity as flies can perform compensatory feeding. However, flies are non-continuous feeders that normally feed little during the night [58,66,67]. During prolonged nocturnal rest phases, the loss of AKH-dependent carbohydrate mobilization leads to increasing AKH release, which in turn positively modulates OANs and subsequently leads to the observed increase in late nocturnal activity, and earlier and more pronounced onset of morning feeding activity in APC-ablated flies kept on food. This scenario is in line with the previous finding that loss of AKHR induces upregulation of *Akh* expression [8]. It also fits with our observation of a drop in glucose levels in the middle of the night, followed by an increase at the end of the night coinciding with the earlier onset of feeding activity in flies with ablated APCs. These flies obviously compensate the loss of carbohydrate mobilisation by feeding during the nocturnal phase, but without triggering increased nocturnal locomotor activity as also AKH:OAN signalling is impaired. A time-restriced increased feeding in AKH-deficient flies is also suggested by the transcriptional up-regulation of the orexigenic peptides NPF and CCHa2 in AKH mutant flies, which paradoxically was not found to be coupled to increased total feeding [71]. Wildtype flies increase *Akh* mRNA levels 3h after the dark-light or light-dark transition, the time points at which starved flies showed pronounced starvation-induced hyperactivity [68]. Interestingly, times of increased sucrose feeding in flies with ablated APCs occurred around ZT3 (significant increase, coupled with low glucose titres) and ZT15 (small and insignificant increase, coupled with high glucose titres) in our study.

When AKH:OAN signalling is specifically impaired under *ad libitum* feeding conditions, flies are still sufficiently able to mobilise their energy stores (via AKH:fat body signalling) to meet the metabolic requirements during rest at night, as evidenced by the observed normal nocturnal activity patterns. However, when AKH:OAN signalling is impaired during the light phase, the increased energy demand during diurnal physical activity triggers AKH release which is then no longer able to stimulate activity-promoting OA signalling, explaining the observed reduced locomotor activity in *Tdc2>AkhR-RNAi* flies.

AKH has been shown to directly trigger strong calcium responses in AKHR-expressing interoceptive SEZ (subesophageal zone) neurons (ISNs) involved in regulating the homeostatic drive to eat and drink [69]. Thus, the observed effects on locomotor activity through the manipulation of AKH signaling in our study might not only be related to foraging activity, but also to the search of water, supporting the idea of a more general effect of AKH on locomotor activity.

The observed effects of manipulating the two AKH signalling branches are not very strong, and our study does not suggest that AKH is a major factor in shaping circadian locomotor activity patterns. Yet, AKH appears to be one of presumably several factors that fine-tune locomotor activity patterns to meet physiological requirements under dietary stress-free conditions. OANs and APCs receive a multitude of converging and interconnected inputs, especially from insulin-producing cells (IPCs), with extensive feedbacks (see [70–72]). For example, OANs in the SEZ express receptors for AKH and DILPs, and integrate the nutritional status by responding to AKH and DILPs in an antagonistic fashion [9]. In turn, APCs express a range of OA receptors [72]. In addition, APCs are negatively modulated by insulin-producing cells (IPCs), suggesting feedforward signalling from IPCs to OANs via AKH signalling in parallel to direct IPC:OAN signalling [72,73]. AKH signalling is also required for the carbohydrate-dependent release of DILP3 [74], and represses expression of DILP2,3 and 5 [71]. Moreover, feeding and diet as well as gustatory perception also influence activity and sleep [73–75], and AKHR is expressed by gustatory neurons including those that express the sugar-sensing receptor Gr5a [16]. AKH production or release is also modulated by diurnal myokine (unpaired 2)-secretion from muscles [76]. Thus, any manipulation of AKH or OA signalling likely results in complex neuroendocrine changes. This complexity may also underlie differences in the effect and effect size observed between different studies. It has to be noted that 4% sucrose served as food source during the locomotor activity recordings in our study. While this is standard for circadian research and allows flies a long-life span, it represents a state of dietary restriction. Future research has to show how the observed locomotor phenotypes manifest on more complete food sources. At least, the locomotor activity-promoting effect of OA seems not to be mediated by DILP-signalling [44]. DILP-signalling seems to rather function upstream and inhibits AKH and OA signalling as DILP2-3,5 mutants show increased day-time activity dependent on AKH and OA signalling [77]. These findings strengthen the view that AKH and AKH:OAN signalling are directly important to balance activity/rest modulation during day and night phases. It remains to be seen to which extend the observed locomotor phenotype following genetic manipulations can be reproduced by endogenous changes in AKH signaling on food.

### OA and AKH signalling is intertwined

A subset of OANs in the SEZ balance sleep and courtship in male flies. Activation of these octopaminergic MS1 neurons suppressed sleep, while silencing the same neurons led to decreased female-induced sleep loss and impaired mating behaviour in male flies [78]. Furthermore, other OANs in the SEZ are important to integrate AKH and DILP signals to modulate activity levels in response to the nutritional situation [9]. The octopaminergic ASM cluster located in the supraesophageal zone has been identified to further include wake-promoting OANs [79].

Consistent with findings from the cockroach [30,31], we now provide evidence in support of OANs in the VNC to be part of an AKH signalling pathway that modulates activity levels under *ad libitum* feeding conditions. *Tdc2>AkhR-RNAi* flies showed reduced activity levels during day-time, while this phenotype was reduced in flies expressing *tshGal80*, which blocks expression in OANs located in the VNC. However, around 7 Tdc2Ns in the VNC were still labelled despite the expression of *tshGal80*. Future studies will clarify whether the slightly lower day activity displayed by *Tdc2,tshGal80>AkhR-RNAi* flies is based on those persisting cells in the VNC or due to OANs in the central brain normally expressing AKHR. An involvement of OANs in the VNC in AKH signalling is further suggested by the observed slow but quite strong increase in intracellular cAMP in mesothoracic OANs upon bath-application of AKH onto dissected VNCs. The slow cAMP increase might suggest indirect action, yet transgenic (over)expression of AKHR in the OANs only mildly shortened the lag time from 15 to around 10 min. Interestingly, the AKH-induced increase in intracellular Ca^2+^ activity of larval prothoracic gland cells sets in only after 15 min of peptide bath application [80]. Clearly, further studies addressing AKHR expression in OANs and the time course of AKH signalling are required to clarify whether the observed cAMP increase in OANs is due to direct or indirect action of AKH. An attempt to address AKHR expression in OANs by available intragenic Gal4/LexA lines did not provide a conclusive answer in our hands (data not shown). In support of an evolutionary conserved activity-promoting action of AKH in the VNC, injection of AKH into the mesothoracic neuropil of the moth *Manduca sexta* leads to increased activity of motoneurons [81].

Our results provide several lines of evidence for a modulation of APC activity by OA and the OAMBII-K3 receptor. A recent cell-specific transcriptomic study supports the expression of OAMB in APCs, yet found expression of further OA receptors (Octbeta1-3R, Oct-TyrR; [72]). At the moment, we can only speculate about the functional significance of OA modulation of APCs and the role of the different OA receptors therein.

## 6. Material and Methods

### 6.1 Fly strains

Flies were cultured according to standard methods. Vials were kept under constant conditions at 25°C and 60% humidity in a 12:12 light:dark cycle. Genetic controls were obtained by crossing Gal4-driver and UAS-effector lines to *w^1118^*. Transgenic and mutant fly lines used in this study are listed in supplementary table S13.

### 6.2 Behavioural assays

#### 6.2.1 Locomotor activity recording

3-7d old flies were recorded individually in *Drosophila* activity monitors (DAM, Trikinetics, Waltham, USA) under standard 12:12 light:dark conditions with light intensities around 60-100lux at ~20°C. Flies were kept in small glass tubes with 2% agarose and 4% sugar on one side. Under starved conditions, flies were kept on 2% agarose to avoid dehydration. 32 flies of a genotype (half male and female) and appropriate controls were measured simultaneously. Fly activity was defined by the number of infrared light beam crosses within 12 hours. Activity patterns were separated into day (ZT0-ZT12; ZT0=lights-on, ZT12=lights-off) and night (ZT12-ZT24), and data was pooled from six days and nights, respectively. For average activity profiles, fly activity was defined as beam crosses per hour. Within these average activity profiles, asterisks label those time points at which experimental flies differ significantly from both genetic controls. For thermogenetic inhibition or activation via *UAS-shibire^ts^* and *UAS-dTrpA1*, experiments were performed at ~31°C under the same light regime.

#### 6.2.2 Survival

Survival analysis was performed with the same DAM system raw data as above. The number of flies alive at the end of day was counted based on present or absent beam crossings.

#### 6.2.3 Startle-induced negative geotaxis (SING)

Negative geotaxis was performed to screen for general motor impairments. The experiments were performed as described in Ali et al. 2011, with minor modifications: Groups of 10 male and female flies, respectively, were separated into Falcon tubes 5 minutes before the experiment. Flies were tapped to the conic bottom of the Falcon tube and then recorded with a DMK22BUC09 video camera with a Pentax C2514-M objective (The Imaging Source Europe GmbH, Germany). Data was averaged over 5 trials per group (with an interspace period of 1 minute) and represents the percentage of flies reaching a label 8.5 cm above the bottom within 5 and 10 seconds.

#### 6.2.4 Flypad feeding assay

Feeding was automatically monitored using a 2-channel flyPAD system (www.flypad.rocks) adapted for long-term measurement of food intake. The flyPAD relies on capacitative measurements [60]. For each experimental run, four males and four females of each genotype were in parallel anesthesized on ice, and then transferred to individual arenas in the late afternoon prior to the evening activity phase.

In the arenas, flies had the choice between 2% agarose (Carl Roth, Karlsruhe, Germany) containing 5 mM sucrose (molecular biology grade, Carl Roth, Karlsruhe, Germany, “sucrose”) or 10% yeast extract (AppliChem, Darmstadt, Germany, “yeast”). Food was provided in glass tubes with an enlarged diameter of 7mm to allow for long-term monitoring. The flyPADs were turned upside-down and placed on evenly-illuminated white LED plates under control of a timer to generate a light regime of LD12:12 in a dark box at 21°C. Feeding (number of sips) was analysed in 30 min bins under exclusion of flies that did not feed during the entire experiment. The data was analysed using custom made Matlab scripts [60].

### 6.3 Live cell imaging

#### 6.3.1 Imaging setup

For Ca^2+^ imaging, specimens were imaged with an AXIO Examiner D1 upright microscope (Carl Zeiss, Göttingen, Germany) with a Zeiss W Plan-Apochromat 40x 1.0 or 20x 0.8 water immersion objective and a SPECTRA 4 hybrid solid state LED source (Lumencor, Beaverton, USA). Images were captured with a PCO.edge 4.2m sCMOS camera (PCO AG, Kelheim, Germany). For cAMP imaging, we used an Axioskop 2FS plus microscope (Carl Zeiss, Göttingen, Germany) with a Visichrom high-speed polychromator system and a Xenon arc lamp (75W, Visitron Systems, Puchheim, Germany). Images were recorded with a CCD camera (CoolSnap HQ, Roper Scientific, Tuscon, USA) and a Photometrics DualView2 beam splitter allowing to separately record CFP and YFP emissions. Images were analysed using VisiView3.0 software (Visitron Systems, Puchheim, Germany).

#### 6.3.2 Bath application experiments

Intracellular Ca^2+^ and cAMP levels were monitored by UAS-*Epac1-camps50A* [48] and UAS-*GCaMP6m* [49]. 1-2 brains were dissected, mounted in 405μl haemolymph-like HL3.1 saline solution [50] in the imaging chamber and allowed to settle for 30-45 minutes. For Ca^2+^ and cAMP imaging in OANs, AKH (+ 0.1% DMSO) or HL3.1 (+0.1% DMSO; as vehicle control), were bath-applied after 100s. Application of carbamylcholine (carbachol; for Ca^2+^ imaging; Fig.S6) and the water-soluble forskolin derivate NKH477 served as positive control (data not shown).

Prior to Ca^2+^ imaging in AKH cells, Ca^2+^ levels were reduced to background levels by applying trehalose (80mM) after 100 seconds [6]. 80mM trehalose was added to all subsequent solutions applied during the experiments. At 300 seconds, either HL3.1 (+0.1% DMSO) as vehicle control or 1mM OA dissolved in HL3.1 (+0.1% DMSO) was bath-applied. Application of potassium chloride (end concentration: 80 mM) after 2980 seconds served as viability control.

Changes in the GCaMP fluorescence intensity within the region of interest were calculated as: ΔF/F_0_=(F_n_-F_0_)/F_0_, where F_n_ is the fluorescence at time point n and F0 the baseline fluorescence intensity calculated from the first 30 seconds. For Epac1 imaging, the CFP/YFP ratio was calculated for each time point: Δ(CFP/YFP) with CFP=CFP_n_/CFP_0_ and YFP=(YFP_n_-(CFP_n_*spillover into YFP)/YFP_0_, where CFPn or YFPn is the fluorescence intensity at a given time point, and CFP_0_ or YFP_0_ is the baseline fluorescence intensity calculated from the first 30 seconds. CFP spillover was determined as 35.7% of the YFP signal.

Region of interests (ROI) were drawn over the cell bodies and the background was subtracted for every ROI. All imaging experiments were performed during the light phase.

### 6.4 Immunostainings

#### 6.4.1 Whole mounts

Immunofluorescence stainings of *Tdc2>10xmyr::GFP* and *Tdc2,tshGal80>10xmyr::GFP* preparations were performed as described [82]. In short, ventral nerve cords of 3-7 days old flies were dissected in HL3.1 (pH 7.2-7.4). Afterwards, specimens were fixated in 4% paraformaldehyde in PBS (0.1M; phosphate buffer saline) for two hours, washed four times in PBS with 0.3% Triton-X 100 (PBT), and blocked with 5% normal goat serum in PBT. Then, specimens were incubated with rabbit anti-GFP serum (A6455, Invitrogen, USA; 1:1000) and mouse monoclonal antibodies against Synapsin (3C11; [83]; 1:50) in blocking solution for one night at 4°C. On day two, preparations were washed six times with PBT and incubated for one night at 4°C with secondary antibodies goat anti-rabbit Alexa 488 (Invitrogen, USA; 1:250) and goat anti-mouse DyLight 649 (Jackson ImmunoResearch, USA; 1:250). Finally, specimens were rinsed six times in PBT and mounted in 80% glycerol in PBS. Until scanning with a Leica TCS SPE or SP8 confocal microscope (Leica microsystems, Wetzlar, Germany), specimens were stored in darkness at 4°C.

#### 6.4.2 Tissue sections

To visualize the arborisations of OANs on the CCA-HG complex, whole body sections were performed (as described in [22]). In short, whole flies with opened cuticle were fixated with 4% paraformaldehyde in PBS for two hours and afterwards washed three times with PBS. Subsequently flies were embedded in hot 7% low melting agarose. After hardening, specimens were cut with a vibratome into 90-95 μm sections. Staining of the sections was continued as described in 6.4.1 in a 24 well plate. Rabbit anti-Tdc2 (pab0822-p, Covalab, France; 1:200) and mouse anti-Synapsin (3C11; 1:50) were used as primary antibodies visualized by goat anti-rabbit Alexa488 (1:200) and goat anti-mouse DyLight649 (1:200) sera.

### 6.5 CaLexA labelling

For CaLexA labelling, equal numbers of 4-11 day old males and female flies were used. The proventriculus plus attached retrocerebral complex (corpora allata and fused corpora cardiaca-hypocerebral ganglion CCA-HG complex) were dissected in HL3.1 saline as previously described [84] and immediately imaged with an AXIO Examiner D1 upright microscope (Carl Zeiss, Göttingen, Germany) with a Zeiss W Plan-Apochromat 40x 1.0 water immersion objective, and a SPECTRA 4 hybrid solid state LED source with 475 nm excitation maximum (Lumencor, Beaverton, USA). Images were captured with a PCO.edge 4.2m sCMOS camera (PCO AG, Kelheim, Germany) at 16bit intensity resolution. All settings were kept constant throughout the experiments. Images were analysed using Fiji [85]. A region of interest was drawn around the CCA-HG and in the direct vicinity (background). The mean intensity of the CCA-HG was then calculated and the mean background was subtracted. In addition, the number of labeled cells was counted.

### 6.6 PCR

Total RNA was extracted from adult Canton-S flies using the Quick-RNA MicroPrep Kit (Zymo Research, Irvine, USA) according to manufacturer’s instructions. Central brains, VNCs, and HG-CCA (for OAMB 15 flies per group; for AKHR five flies per group) were dissected in HL3.1 medium, and transferred to 300 μl RNA lysis buffer on ice. Tissues were homogenized with a plastic pestle. Total RNA was eluted in 8 μl RNAse-free water. For cDNA synthesis, the QuantiTect Reverse Transcription Kit from Qiagen (Hilden, Germany) was used. All steps were performed following the manufacturer’s protocol. Genomic DNA was removed by adding 1 μl of gDNA wipe-out buffer to 6 μl of the eluted RNA. Following incubation at 42°C for 2min, the samples were placed for 2min at 4°C and 3 μl of a mastermix composed of 2 μl RT buffer, 0.5 μl RT primer Mix and 0.5 ml reverse transcriptase was added. Reverse transcription was performed for 30 min at 42°C, followed by 3 min at 95°C and 2min at 4°C. Finally, 20 μl water were added; cDNA samples were stored at −20 °C. To analyse OAMB and AKHR expression in the isolated tissues, cDNAs were PCR-amplified using specific primers (Table 1) and a JumpStart REDTaq ReadyMix Reaction Mix (Sigma-Aldrich, MO, USA). α-tubulin and water (“no DNA”) were used as internal control. The PCR programme consisted of 5min at 95 °C, followed by 35 cycles of 30 s at 95°C, 30 s at 61°C (57,5°C for AKHR), 60 s at 72°C and finally 5 min at 72°C.

**Table 1:**
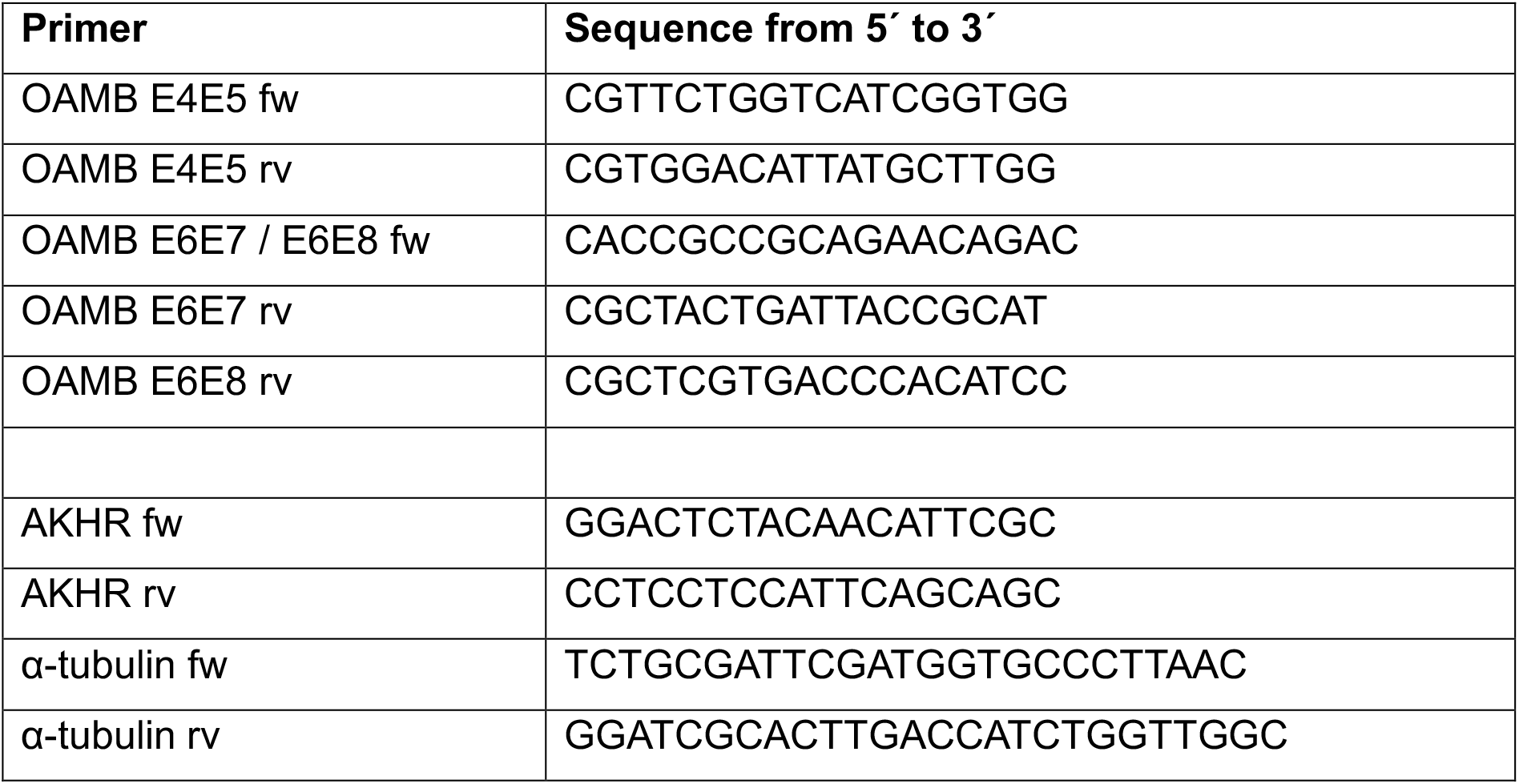
Primer pairs used for PCR in this study.

### 6.7 Ultra performance liquid chromatography-tandem mass spectrometry (UPLC-MS/MS) measurements

#### 6.7.1 Haemolymph sampling

For each sample, 20 flies were perforated by a thin needle at the thorax and put in a perforated 0.5ml Eppendorf cap, which was fitted into a 1.5ml Eppendorf cap. Haemolymph was collected within the 1.5ml Eppendorf cap after 1min of centrifugation in a benchtop centrifuge at 5000 rpm, and immediately deep-frozen until analysis.

#### 6.7.2 UPLC-MS/MS analysis of sugars and diacylglycerols

Sugars were analysed using a Waters Acquity ultra-high-performance liquid chromatograph coupled to a Waters Micromass *Quattro Premier* triple quadrupole mass spectrometer (Milford, MA, USA) with an electrospray interface. Chromatographic separation was performed on a Waters Acquity BEH amide column (1.7 μm, 2.1×100mm; Waters, Milford, MA, USA) according to Waters application note WA60126 with a modified flow rate of 0.2 ml/min. The analytes were eluted in a linear solvent-strength gradient (75% to 45% acetonitrile containing 0.1% ammonium hydroxide in 10min) at a column temperature of 35°C. Sugars were detected in the negative electrospray mode at a source temperature of 120°C and a capillary voltage of 3.25 kV. Nitrogen was used as desolvation and cone gas with flow rates of 800 l h^−1^ at 350°C and 25 l h^−1^. The mass spectrometer was operated in multiple reaction monitoring (MRM) mode using Argon as collision gas at a pressure of approximately 3 x 10^−3^ bar. Cone voltage (CV) and collision energy (CE) were optimized for maximum signal intensity of each individual compound during ionization and collision induced dissociation (CID) for a dwell time of 50ms per transition (Table 2). Glucose-6,6-d2 and Trehalose-1,1’-d_2_ were used as internal standards for quantification of mono- and disaccharides.

**Table 2:** Table with specific cone voltage and collision energy for each transition.

| Compound Name | Parent (m/z) | Daughter (m/z) | CV (V) | CE (eV) |
| --- | --- | --- | --- | --- |
| Glucose/Fructose | 179 | 89 | 18 | 10 |
| Glucose-6,6-d<sub>2</sub> | 181 | 91 | 18 | 10 |
| Sucrose/Trehalose | 341 | 179 | 32 | 15 |
| Trehalose-1,1'-d<sub>2</sub> | 343 | 180 | 32 | 15 |

Determination of diacylglycerols were performed according to [86]. Prior to analysis, haemolymph samples were dried and recovered in 75 μl isopropanol containing 1 μg/ml 1,2-didecanoyl-glycerol used as an internal standard.

### 6.8 Statistics

Data was analysed for normal distribution using the Shapiro-Wilk test. For statistical comparison between genotypes, a pairwise t-test was used for normally distributed data, a pairwise Wilcoxon rank sum test was used for not normally distributed data (both including a Bonferroni-Holm correction for pairwise comparisons). All statistical analyses were done with R Studio version 1.0.136. (www.r-project.org). For the statistical comparison of sugar levels quantified by LC-MS, we used a simple t-test based on the relatively small sample size (N=3-5, with n=20 per N). Data were plotted using OriginPro 2016G, GraphPad Prism 8, R Studio and Photoshop CS6. Activity levels are presented as box plots, with 50% of values for a given genotype being located within the box. Box and whiskers represent the entire set of data. Outliers are indicated as open circles. The median performance index is indicated as a thick line within the box plot. Data presented as barplots show the mean values and standard error of the mean (SEM). Significance levels between genotypes shown in the figures refer to the raw p-values obtained in the statistical tests.

## Acknowledgements

The authors thank Bryon Hughson, Wolf Hütteroth, Robert Kittel and Pamela Menegazzi for fruitful discussions and comments on the manuscript, Gertrud Gramlich and Susanne Klühspies for excellent technical assistance, and Jay Hirsh and the Bloomington Stock centre for providing flies. This work was supported by intramural support of the University of Würzburg (to C.W.) and the University of Graz (to R.P.K.), and the Deutsche Forschungsgemeinschaft (SFB1047 “Insect timing”, project B2 to C.W., DP1979/3-1 to D.P., PA3241/2-1 to M.S., and FO207/14-1 to C.H.F.). The authors declare no competing interests. D.P., M.S., J.R., R.P.K. and C.W. conceived and designed the experiments. D.P., M.S., J.R., K.A., M.K., C.H.-L., A.G.Ü., N.E., P.M.I. and C.W. performed the experiments. D.P., M.S., J.R., A.F., M.K., C.H.-L., P.M.I., C.H.F., M.J.M., and C.W. analysed the results. D.P. and C.W. wrote the article. All authors provided comments and approved the manuscript.

## Supplement Figures

**(S1-S2).**
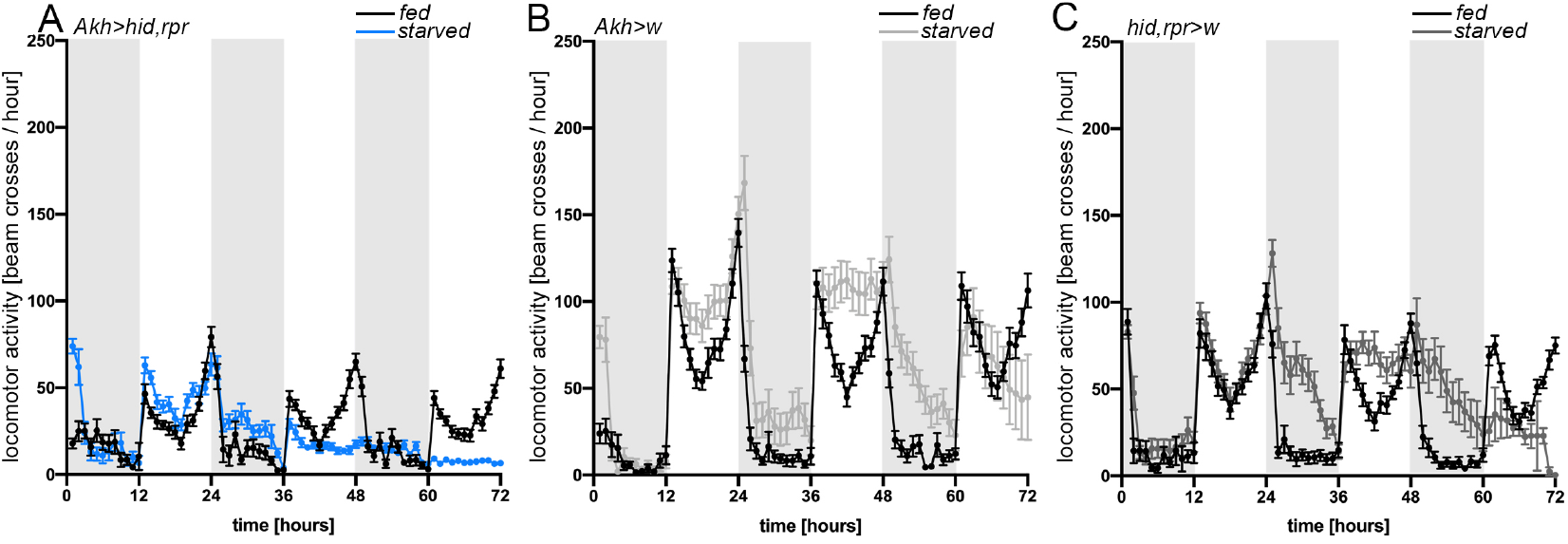
In response to starvation Akh>hid,rpr flies show only a minor increase in activity, while there was no starvation-induced increase in activity within the first 48h in contrast to control flies.

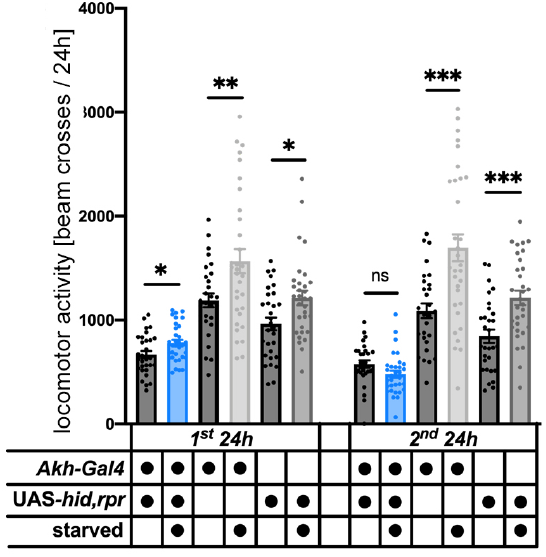

**(S3).**
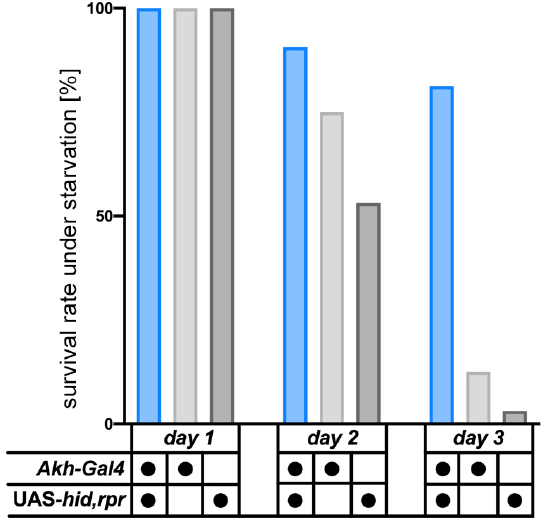
The loss of AKH signalling increased the starvation susceptibility as control flies died earlier compared to Akh>hid,rpr flies.

**(S4).**
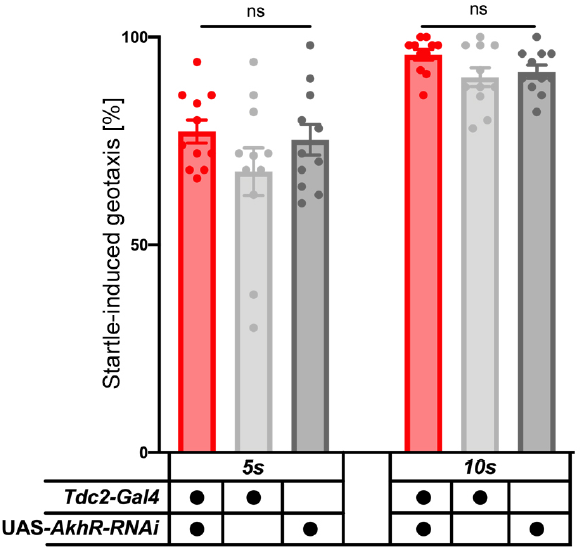
The knockdown of AKHR in OANs did not affect motor behaviours in general, as experimental flies performed indistinguishable from genetic controls in the startle-induced climbing assay (all p>0.05; all N=11).

**(S5).**
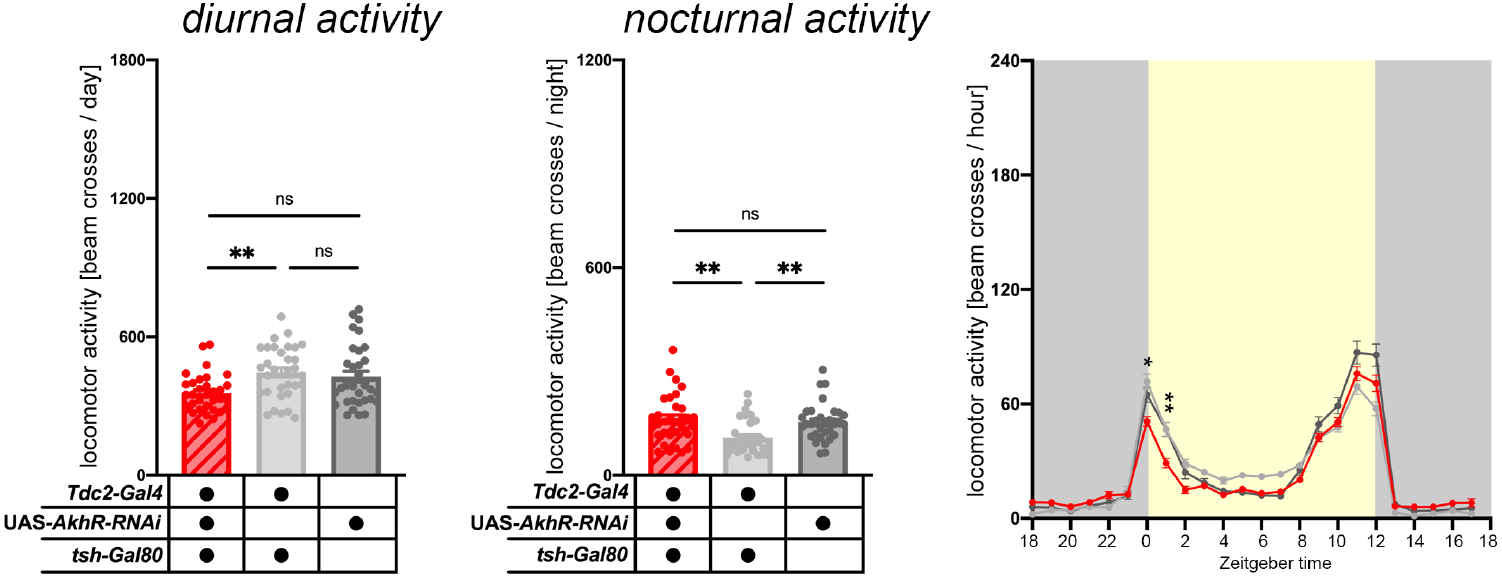
Specific knockdown of AKHR in Tdc2Ns (Tdc2-Gal4-positive neurons) of the VNC specifically affected locomotor activity.

**(S6).**
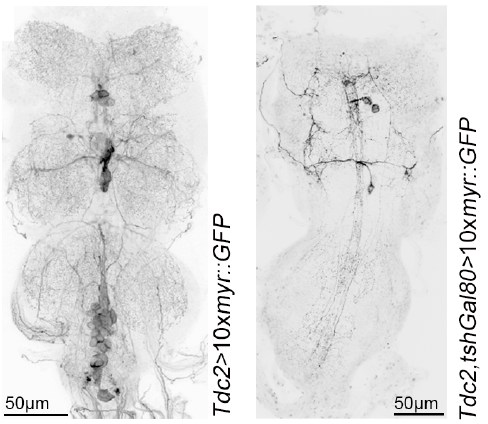
Expression of tshGal80 restricted the expression pattern of Tdc2-Gal4 within the VNC to ~7.4 ± 2.07 cells (n=5

**(S7).**
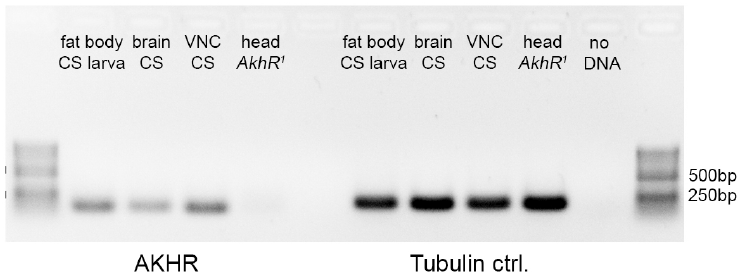
RT-PCR analysis revealed that AKHR is expressed in larval fat bodies, the adult brain and VNC. Significance levels: p>0.05 ns, p<0.05 *, p<0.01 **, p<0.001 ***. [CS= CantonS, VNC=Ventral nerve cord].

**(S8).**
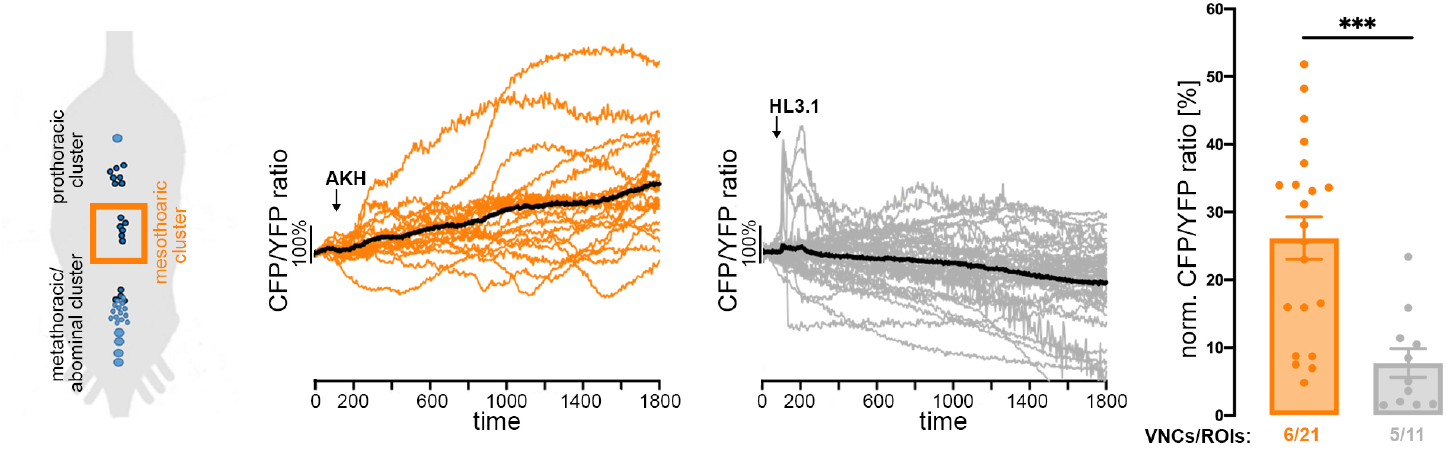
cAMP imaging experiments in Tdc2Ns (Tdc2-Gal4-positive neurons) of the mesothoracic cluster (orange box) in the VNC revealed that cells significantly responded with a cAMP increase to AKH bath application.

**(S9).**
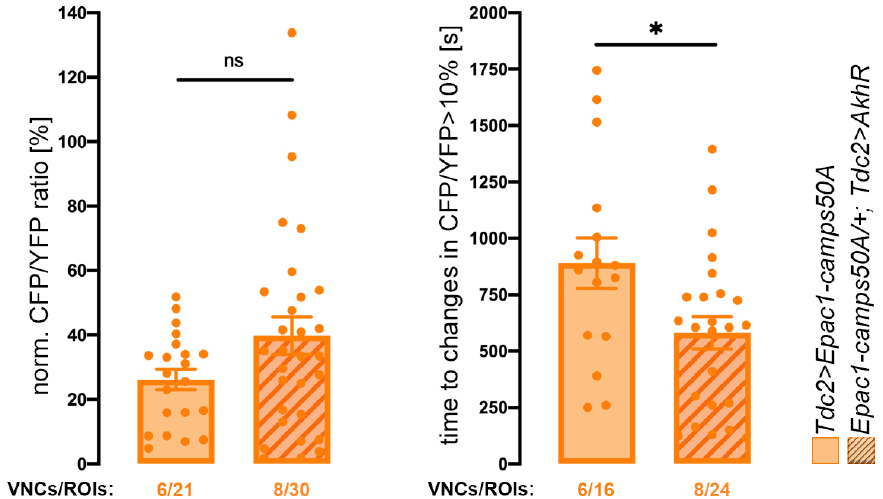
Overexpression of AKHR did not affect response levels and did only slightly reduce the lag time in response to AKH in Tdc2Ns.

**(S10).**
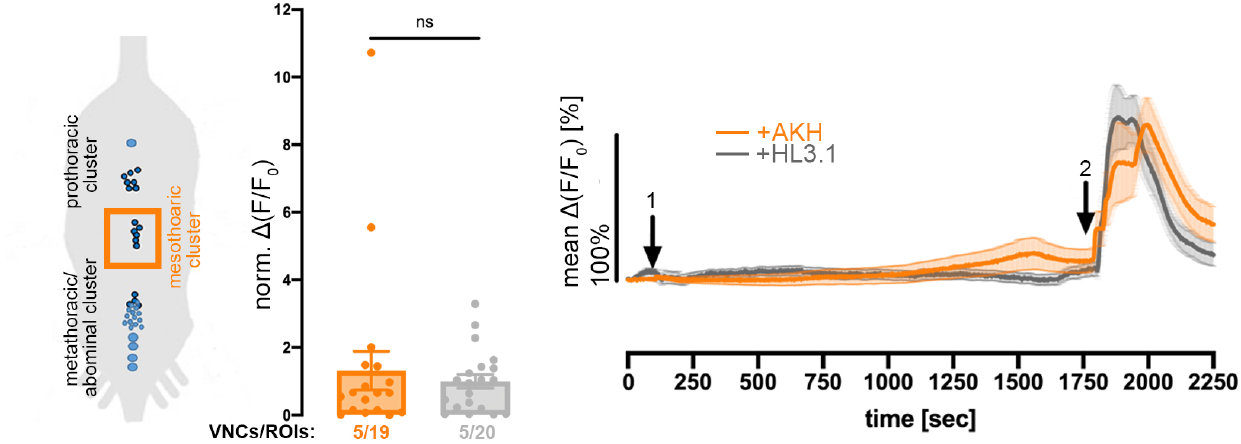
Ca^2+^-imaging in Tdc2Ns of the mesothoracic cluster in the VNC revealed that these cells did not respond with a significant Ca^2+^ increase to AKH bath-application (arrow 1 at 100s). Application of carbamylcholine at 1800s (arrow 2) served as a positive control for viability of the cells. [ROI=Region of interest, VNC=Ventral nerve cord]. Error bars indicate SEM.

**(S11).**
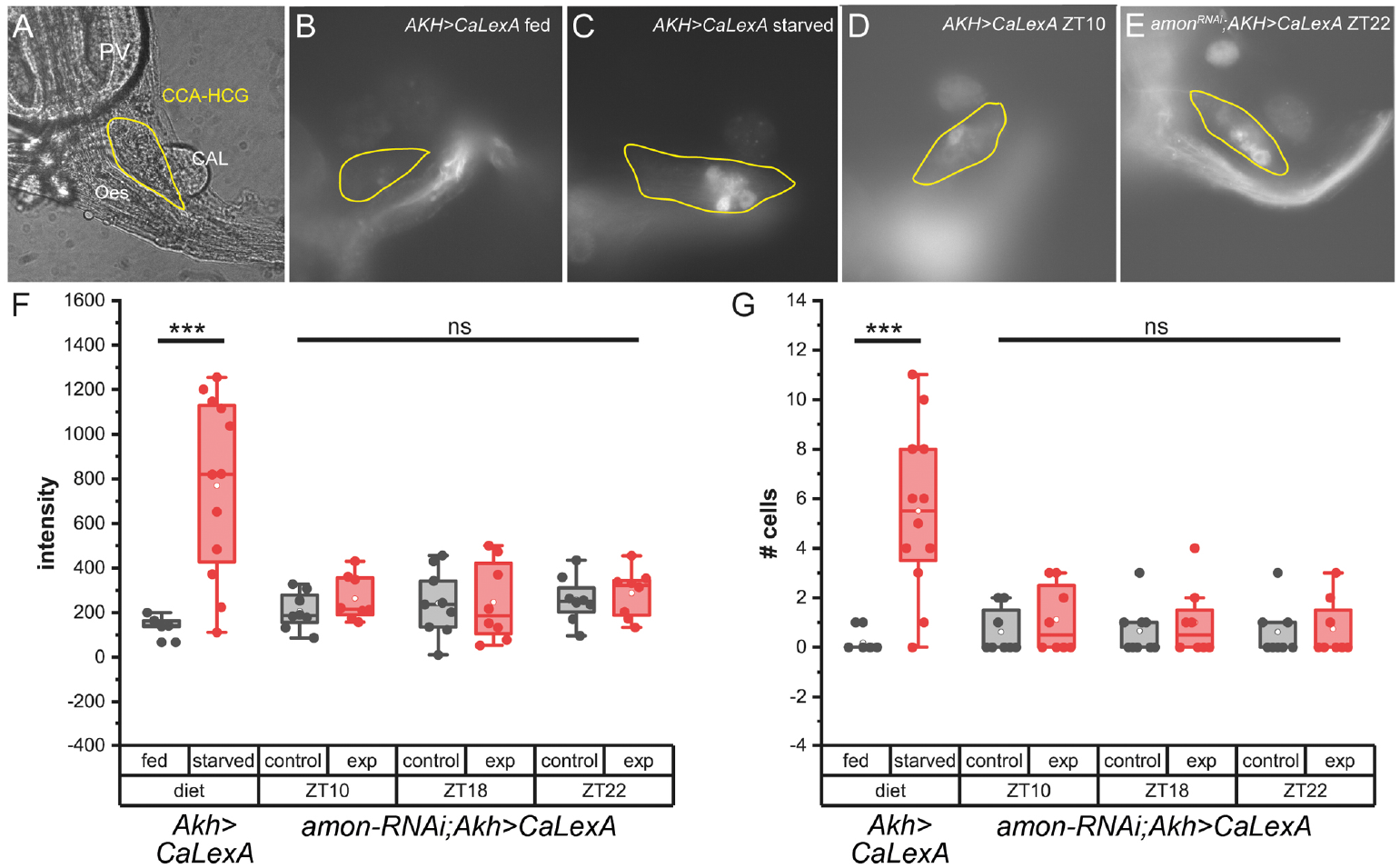
CaLexA-induced GFP expression in AKH-producing cells (APCs). **(A)** shows a typical preparation of the CCA-HG complex including the corpora allata (CA), esophagus (ES) and proventriculus (PV). **(B-E)** Examples for CaLexA-induced GFP expression in Akh>CaLexA flies on food (B) and 24h off food (C), as well as Akh>CaLexA controls at ZT10 (D) and amonRNAi;Akh>CaLexA experimental flies at ZT22 (E). **(F-G)** CaLexA-induced GFP expression in APCs under fed (n=6) or starved (n=12) conditions (Akh>CaLexA, left two columns) and on food at different time points during the day (Akh>CaLexA = controls, amonRNAi; Akh>CaLexA = exp (experimental flies), n=8-9 per condition) quantified as GFP fluorescence intensity in (F), and number of GFP-expressing APCs in (G). Significance levels: p>0.05 ns, p<0.05 *, p<0.01 **, p<0.001 ***. [APCs= AKH-producing cells; ZT= Zeitgeber time].

**(S12).**
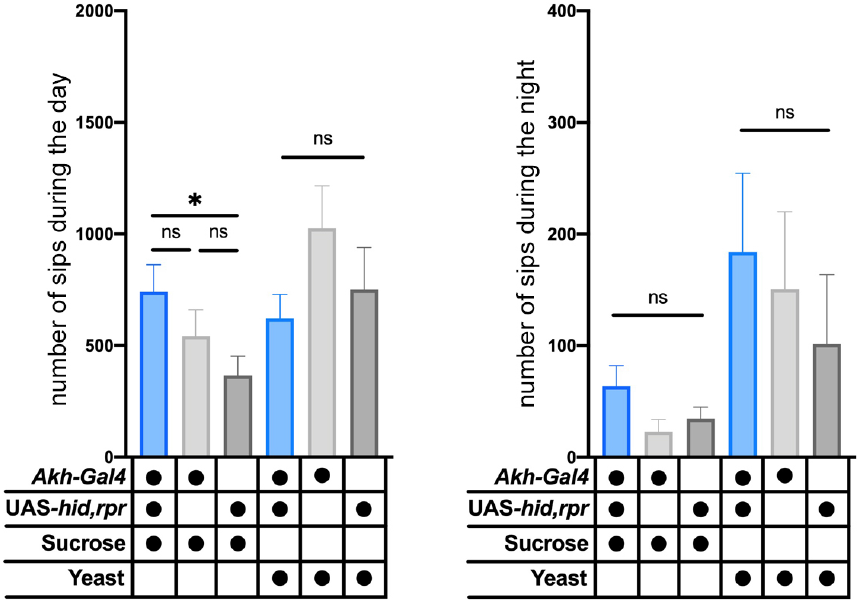
Total feeding indicated by the number of sips was not affected by APCs for both diurnal and nocturnal sucrose and yeast consumption. n=35-36; signifcance levels p<0.05 *, p<0.01 **, p>0.001 ***. Error bars indicate SEM.

**S13.**
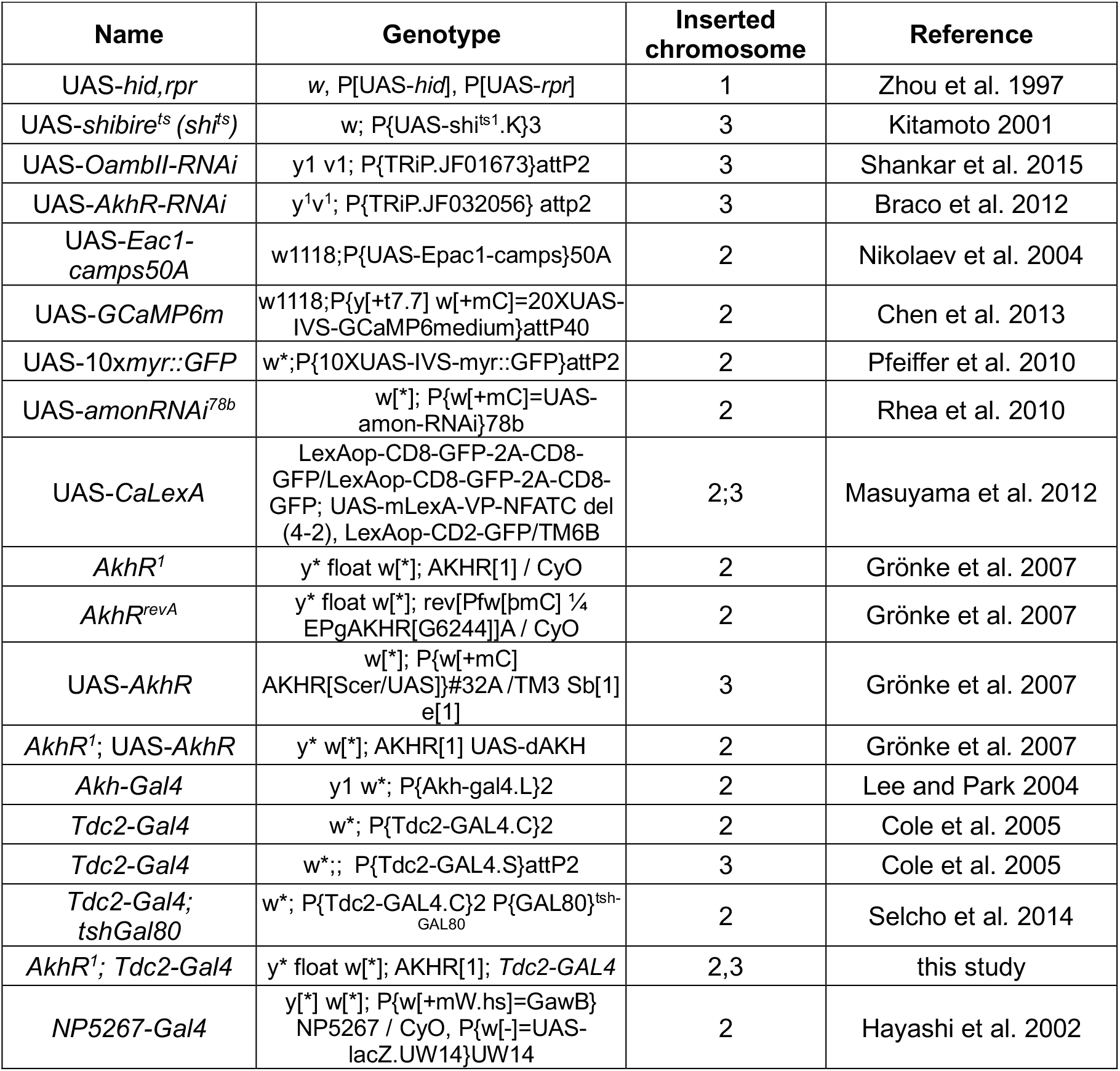
Fly lines

